# Somatic variants activating the RAS-MAPK pathway confer susceptibility to hippocampal sclerosis in drug-resistant epilepsy

**DOI:** 10.64898/2026.04.06.716727

**Authors:** Lauren Mashburn-Warren, Ashton Holub, Sahibjot Sran, Swetha Ramadesikan, Keaton R. Suh, Allison Thompson, James J. Anderson, Adithe Rivaldi, Ari R. Zavarella, Megan Chandler, Allison Daley, Corinne H. Strawser, Elizabeth A. R. Garfinkle, Jonathan Pindrik, Ammar Shaikhouni, Jeffrey Leonard, Daniel R. Boué, Diana L. Thomas, Christopher R. Pierson, Elaine R. Mardis, Katherine E. Miller, Adam P. Ostendorf, Daniel C. Koboldt, Tracy A. Bedrosian

## Abstract

Hippocampal sclerosis is a frequent finding in pediatric epilepsy surgery and has traditionally been regarded as an acquired lesion. It commonly co-occurs with focal cortical dysplasia (FCD IIIa), yet whether hippocampal injury is secondary to seizures or reflects a shared underlying etiology remains unresolved. Here we identified somatic variants activating the RAS–MAPK pathway in 40% of patients with hippocampal sclerosis, but in none with non-sclerotic hippocampus. Gain-of-function variants in *PTPN11* were the most common finding, with mutations present in both cortex and hippocampus and enriched in hippocampal neurons, consistent with a shared developmental origin. In mice, *Ptpn11*^D61Y^ mutants developed profound hippocampal degeneration and gliosis following subthreshold kainic acid exposure, whereas wild-type controls were unaffected. p38-dependent stress pathways were upregulated in patients and mice, suggesting a mechanism through which ERK-p38 crosstalk lowers the threshold for seizure-induced injury. These results provide a genetic explanation for FCD IIIa, elucidate the role of somatic mutations within the RAS-MAPK pathway in driving hippocampal sclerosis, and provide a target for pathway-specific interventions for intractable seizures.

## Main

Malformations of cortical development are a major cause of drug-resistant epilepsy in children (1). Focal cortical dysplasia (FCD) is the most common malformation, which is classified into three types based on histopathological features (2). FCD I is characterized by abnormal radial and/or tangential cortical lamination, FCD II is defined by dyslamination with dysmorphic neurons with or without balloon cells, and FCD III represents cortical dyslamination associated with a principal lesion, including hippocampal sclerosis (IIIa), low-grade tumor (IIIb), vascular malformation (IIIc), or another acquired lesion (IIId) (3). In FCD III, the relationship between abnormal cortical architecture and the associated lesion remains incompletely defined. Specifically, it is unknown whether the two pathologies have distinct causes or arise through a common mechanism.

Over the past decade, brain-restricted somatic genetic variants have been implicated in the pathogenesis of FCD and there appear to be strong genotype-phenotype correlations. For example, variants activating the PI3K-AKT-mTOR pathway are identified in approximately half of individuals with FCD II with the variants enriched in histologically abnormal cell types (3). Genetic associations for FCD I are not as clearly defined but seem to involve somatic variants affecting pathways other than mTOR activation (3–8). There is currently no known genetic cause for FCD IIIa (3). However, somatic variants activating the RAS-MAPK pathway (e.g., *PTPN11, SOS1, KRAS, BRAF,* and *NF1)* are identified in hippocampal tissue from at least 10% of adult patients with mesial temporal lobe epilepsy (MTLE) (9), which is frequently associated with hippocampal sclerosis.

These findings suggest somatic variants activating the RAS-MAPK pathway may also contribute to FCD IIIa, but the relationship between its cortical versus hippocampal components remains unknown. In children, contrary to adults, hippocampal sclerosis is rarely observed as the sole defining pathological feature. Rather, it presents with adjacent cortical dysplasia as FCD IIIa in as many as 79% of cases (10) suggesting the two pathologies result from a common mechanism. However, cortical dysplasia is present from birth, whereas hippocampal sclerosis is rarely observed in children under 2 years of age, suggesting it develops over time (11). This could indicate that hippocampal sclerosis is a secondary consequence of seizures that arise primarily from abnormal cortex, although it’s unknown why some children with FCD develop hippocampal sclerosis and others do not.

Here, we investigated whether somatic variants contribute to hippocampal sclerosis in the context of developmental temporal lobe dysplasia by performing genetic analysis of surgically resected brain tissue from pediatric patients treated for drug-resistant epilepsy. We sequenced hippocampal and adjacent cortical tissue when available to determine the anatomical distribution of contributory somatic variants, thereby elucidating common versus distinct etiologies. Next, we generated a mouse model carrying a neuron-specific *Ptpn11* gain-of-function variant to determine whether somatic variation predisposes to hippocampal sclerosis in the setting of seizure activity. Our results demonstrate that somatic variation of the RAS-MAPK pathway contributes to the pathogenesis of FCD IIIa. We suggest that RAS-MAPK pathway somatic variants may underlie a spectrum of epileptic pathologies depending on developmental timing of the variant, ranging from the more widespread, earlier onset FCD IIIa to the more focal, adult-onset MTLE.

### RAS-MAPK Pathway Somatic Variants are Associated with Hippocampal Sclerosis

Pediatric patients undergoing surgical treatment for drug-resistant epilepsy at Nationwide Children’s Hospital were enrolled in an IRB-approved study for genetic analysis of resected brain tissue. In retrospective review, we included as ‘cases’ all patients with a diagnosis of hippocampal sclerosis or mesial temporal sclerosis based on neuropathology and/or neuroimaging findings (Fig. 1A). Most cases had both hippocampal and cortical tissue available for study. Patients who had hippocampal tissue resected but no findings of sclerosis (including diagnoses of FCD I, FCD II, and FCD III) were included as controls (Supplementary Table 1). Each patient’s clinical history was evaluated for risk factors of hippocampal sclerosis, including early-life brain injury, febrile seizures, or status epilepticus. Approximately 60% of cases had at least one of these risk factors compared to 40% of controls (Fig. 1B).

**Figure 1:**
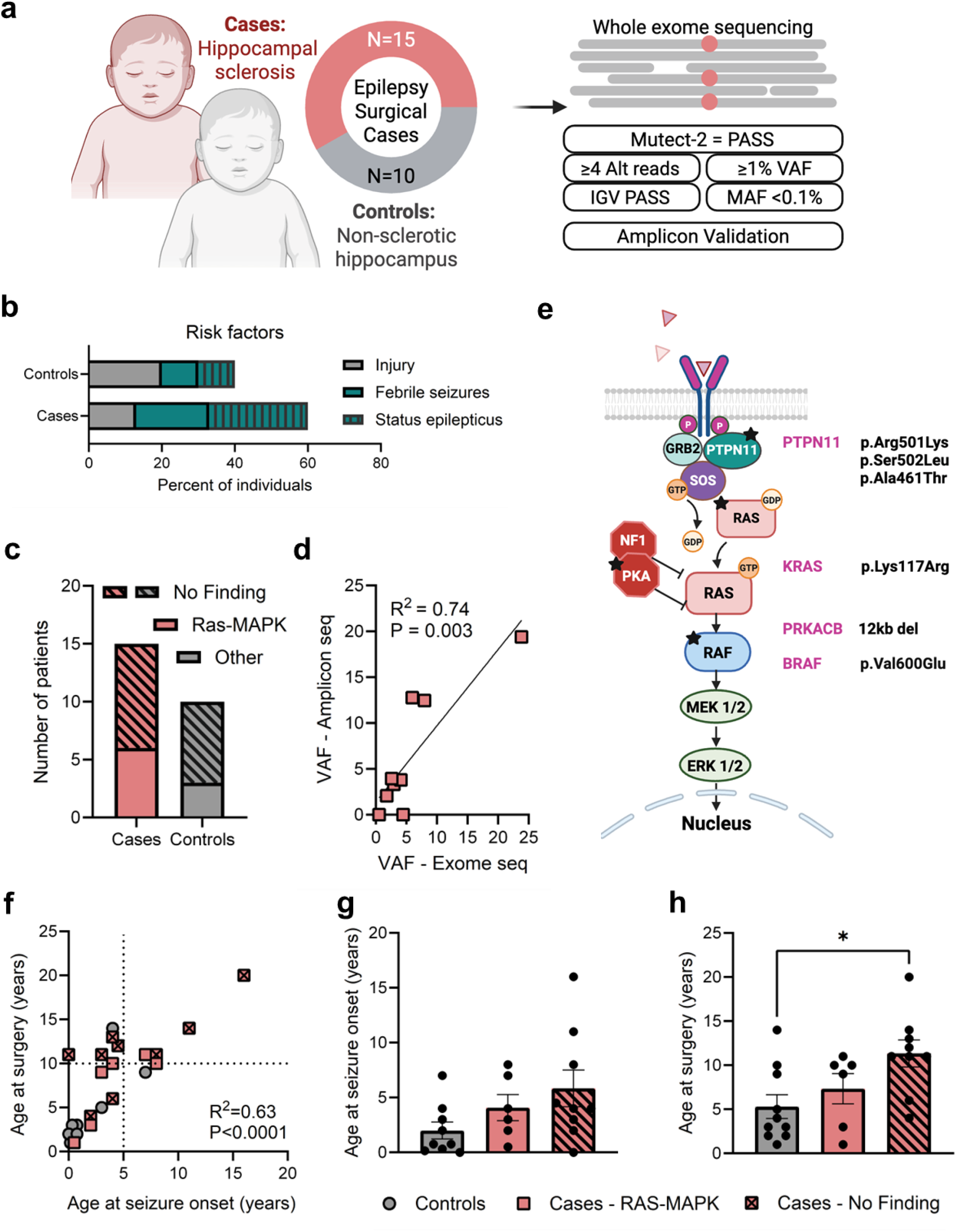
RAS-MAPK Pathway Somatic Variants are Associated with Hippocampal Sclerosis. Surgically resected brain tissue was collected from pediatric epilepsy surgery patients with FCD and hippocampal sclerosis (cases, N=15) or FCD and non-sclerotic hippocampus (controls, N=10) for detection of somatic variants **(A)**. Approximately 60% of cases had a risk factor for hippocampal sclerosis including history of brain injury, febrile seizures, or status epilepticus **(B)**. Exome sequencing analysis identified potentially contributory somatic variants in 6/15 cases, all of which are predicted to activate the RAS-MAPK signaling pathway **(C)**. Somatic variants and variant allele frequencies were verified by amplicon sequencing **(D)**. Schematic showing the variants identified in cases and their role in MAPK signaling **(E)**. Age at surgery was significantly correlated with age at seizure onset among cases and controls **(F)**. Seizure onset age was similar across controls, cases with RAS-MAPK somatic variants, and cases without findings **(G)**. Cases without findings had significantly greater age at surgery. One-way ANOVA; F(2,22)=4.58, P=0.02, Tukey’s post-hoc P=0.02 **(H)**.

For each patient, genomic DNA was extracted from available resected tissue and subjected to whole exome sequencing and analysis in comparison to a matched blood sample to identify somatic genetic variants. Variant call sets were filtered and assessed (see Methods). We detected 9 total candidate somatic variants among cases and controls (Supplementary Table 2). Among cases, 6/6 variants were identified in genes belonging to the RAS-MAPK pathway (Fig. 1C). The remaining three variants identified in the control group belonged to unrelated pathways. All variants were verified by amplicon sequencing, which confirmed variant allele fractions (VAFs) as high as 20% (Fig. 1D). In a case/control categorical comparison, we found RAS-MAPK pathway variants were enriched in cases with hippocampal sclerosis (6/15 cases vs. 0/10 controls, Fisher’s exact test, P=0.05). Narrowing strictly to cases diagnosed as FCD IIIa, and excluding one case of dual pathology, this enrichment was more evident (4/7 FCD IIIa cases vs. 0/10 controls, Fisher’s exact test, P=0.01).

The gene most implicated in cases was *PTPN11* (3/6 variants), which has been previously reported in FCD and mesial temporal lobe epilepsy (4, 6, 9) (Fig. 1E). All three variants that we identified have been previously reported as Pathogenic for RASopathy/Noonan syndrome in the germline context according to the ClinVar database (https://www.ncbi.nlm.nih.gov/clinvar/). *PTPN11* encodes the SHP2 protein for which gain-of-function variants activate downstream MAPK signaling (12). It has also been implicated in cortical development in mouse models (13, 14). In addition, we identified one *KRAS* somatic variant not previously reported; however, other missense changes at this residue have been reported as Pathogenic for various tumors in ClinVar. We also identified the well-known BRAF V600E variant in a patient with hippocampal sclerosis and a low-grade glioneuronal lesion (diagnosed as dual pathology: FCD IIIa / FCD IIIb). Lastly, we found an 11.9 kb deletion of exons 2-5 in *PRKACB*, which encodes the beta catalytic subunit of Protein Kinase A (PKA). Loss of PKA-dependent regulation increases downstream RAS-MAPK signaling in a cell type-dependent manner (15).

No relevant somatic variants were identified by exome sequencing in the nine remaining cases with hippocampal sclerosis; however, there was heterogeneity among the cohort in terms of clinical history, age at seizure onset, and age at surgery that could explain these results. In general, there was a strong linear correlation between age at seizure onset and surgery, however a subgroup of cases with hippocampal sclerosis had >5 years between seizure onset and surgical intervention (Fig. 1F). Age at seizure onset was not significantly different across cases or controls (Fig. 1G); however, age at surgery was increased among cases with no genetic finding (Fig. 1H). This raises the possibility that there are distinct clinically appreciable subgroups of patients with different etiologies. Indeed, cases with a history of encephalopathic hemispheric injury had >5 years between seizure onset and surgical intervention (see Supp. Table 1). Motivated by this heterogeneity in the cohort, we performed an additional case/control comparison excluding individuals with history of perinatal stroke, traumatic brain injury, encephalitis, or tumor. In this restricted ‘developmental’ cohort, we found strong enrichment of RAS-MAPK variants in cases (5/10 cases vs. 0/10 controls, Fisher’s exact test, P=0.03), suggesting these variants are not just a marker of severely injured brain, but rather they may explain a distinct subset of patients with hippocampal sclerosis and developmental cortical dysplasia.

### Somatic Variants Activate RAS-MAPK Signaling

To investigate the tertiary-level interactions of variant residues in patient tissue samples, we used structural modeling to generate folded protein structures and investigated predicted hydrogen bonds formed by these residues within 4Å. Protein structural modeling of the *PTPN11* (SHP2; A461T, R501K, S502L) and *KRAS* (K117R) variants demonstrated architectural changes in the SHP2 catalytic pocket and the KRAS GTP binding site that further suggest pathogenicity.

For wildtype PTPN11, we modeled the protein-tyrosine phosphatase domain and found that A461 colocalizes with R501, and S502 within the tertiary structure in proximity to the catalytic pocket housing the catalytic cysteine, C459 (Fig. 2A). Substitution of these residues with the patient variants is predicted to significantly alter the architecture of the catalytic pocket (Fig. 2B, Supplementary Text). In past studies, functional studies of the A461T mutation showed that it reduces phosphatase activity but simultaneously weakens autoinhibition and enhances ligand-induced activation, resulting in increased downstream MAPK signaling (16). Because R501 and S502 are similarly positioned to A461 near the catalytic pocket of the tertiary SHP2 structure, variants at these residues may have similar effects (17, 18).

**Figure 2:**
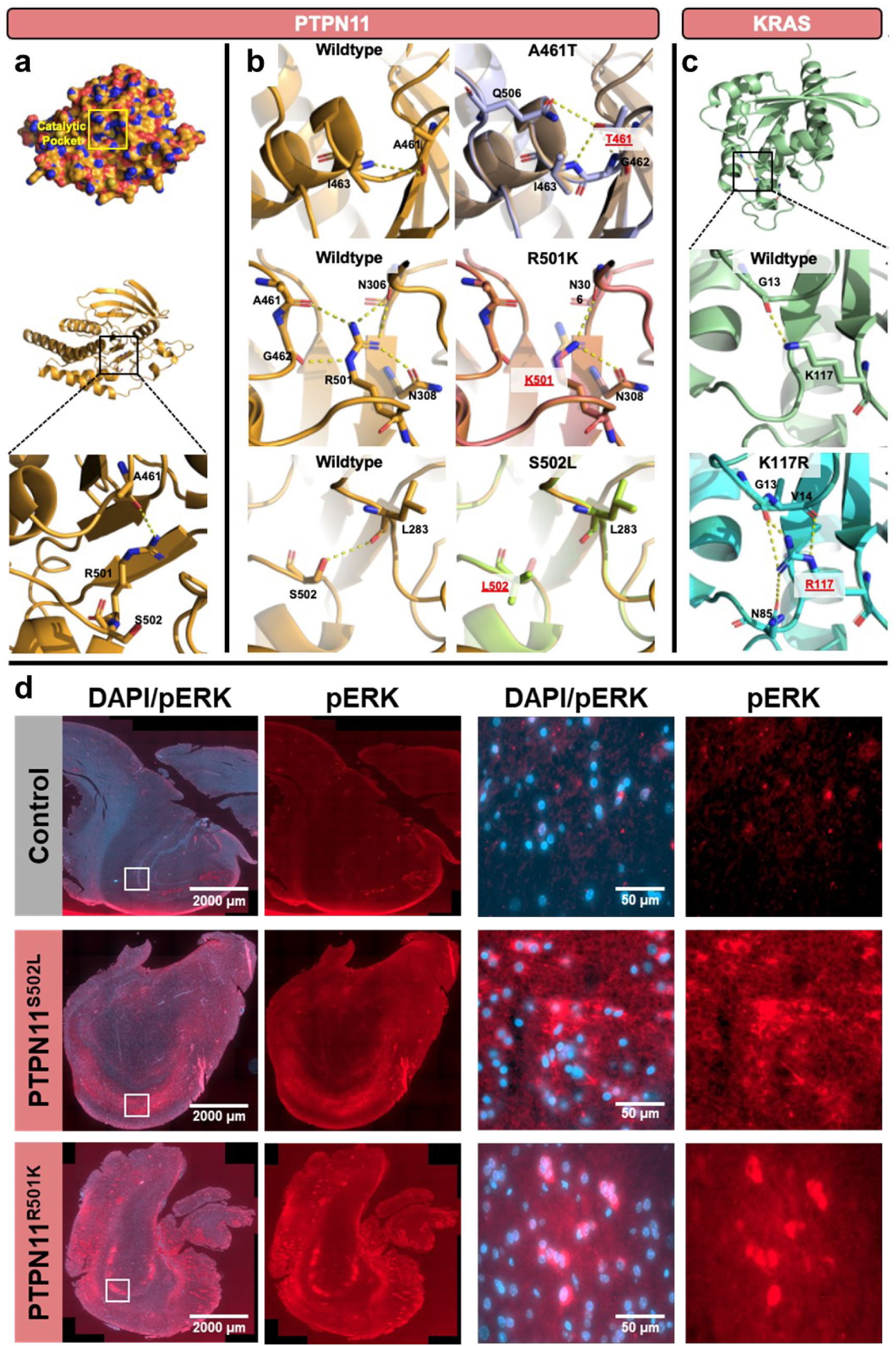
Somatic Variants Activate RAS-MAPK Signaling. Surface view of the catalytic pocket of the PTPN11 protein-tyrosine phosphatase domain and cartoon model showing colocalization of three patient-derived PTPN11 variants: A461T, R501K, and S502L (**A**). PTPN11 patient variants change the architecture of the catalytic pocket with predicted functional effects. The wildtype protein-tyrosine phosphatase domain is orange, and PTPN11 structures carrying the A461T, R501K, and S502L are light blue, pink, and bright green respectively **(B)**. Structural model of KRAS showing location of K117 within the wildtype structure. KRAS K117R forms a stronger interaction with G13 and new interactions with V14 and N85. The wildtype KRAS structure is pale green and the KRAS structure carrying the K117R mutation is teal. Dashed yellow lines indicate predicted hydrogen bonding (≤4.0Å) **(C)**. Immunofluorescence staining of patient hippocampal tissue shows intense phospho-ERK staining in the hippocampus of patients with RAS-MAPK pathway variants, but absent from controls without hippocampal sclerosis **(D)**.

We then investigated the effects of the KRAS K117R mutation and found that the sidechain of K117 within the wildtype KRAS structure is predicted to form an interaction with the backbone G13 (2.8Å), a residue within the GTP binding site which is frequently mutated in cancer (19–21). R117K is predicted to generate a stronger interaction between the sidechain of R117 and backbone of G13 (2.1Å) compared to lysine, as well as a second weaker interaction (3.7Å). Additionally, the sidechain of R117 is predicted to establish new interactions with the backbone of V14 (3.8Å) and sidechain of N85 (3.9Å) (Fig. 2C). In functional studies, K117R as well as G13D mutations have been characterized as having higher rates of guanine nucleotide exchange (22–24).

The expected outcome of these variants is activation of MAPK signaling, culminating in increased levels of phosphorylated ERK (pERK). To confirm the functional effect of these structural changes at the protein expression level, we performed immunofluorescence staining of pERK in patient hippocampal tissue. Cases with RAS-MAPK pathway somatic variants had dramatically increased expression of pERK compared to controls (Fig. 2D).

### Chronic MAPK Signaling Promotes Compensatory Cellular Stress

To investigate downstream transcriptional changes associated with RAS-MAPK pathway somatic variants, we performed bulk RNA-sequencing of fifteen resected samples from the six patients with RAS-MAPK variants and compared gene expression against twelve tissue samples derived from five age- and brain region-matched post-mortem neurotypical donors. Patient samples had markedly distinct transcriptional signatures that clustered separately from donor samples (Fig. 3A). Differential gene expression analysis identified over 600 significant genes, with the majority being upregulated (Fig. 3B and Supp. Table 3). Gene set enrichment analysis identified biological pathways involved in inflammation, KRAS signaling, MTORC1 signaling, and apoptosis (Fig. 3C). Significant upregulation of immediate early genes, apoptosis genes, MAPK downstream effectors, and MAPK feedback regulators was observed in patient tissues (Fig. 3D). Specifically, patient tissues showed induction of the p38–MK2/MK3 stress response axis and selective MAPK feedback regulators (DUSP1, DUSP9, DUSP16), coupled with suppression of the ERK-associated phosphatase DUSP4. Concurrent upregulation of GADD45B and TP53BP2 suggests engagement of stress-adaptive checkpoint programs rather than a purely mitogenic ERK transcriptional state (25–30). These results suggest that chronic activation of RAS signaling elicits compensatory stress and feedback programs in patient tissues at the cost of persistent cellular stress and inflammation. To explore gene expression changes in cases with no genetic finding, we performed a separate comparison against neurotypical control tissue (Ext. Data. Fig. 1A). Interestingly, there were both unique and shared genes with those identified in the cases with RAS-MAPK pathway somatic variants (Ext. Data. Fig. 1B and Supp. Table 4). Gene set enrichment analysis identified overlapping biological pathways to those found in cases with RAS-MAPK variants (Ext. Data. Fig. 1C), but gene ontology analysis of mutual versus shared differentially expressed genes also revealed some unique terms, again suggesting that there are distinct patient subsets within the cohort (Ext. Data. Fig. 1D).

**Figure 3:**
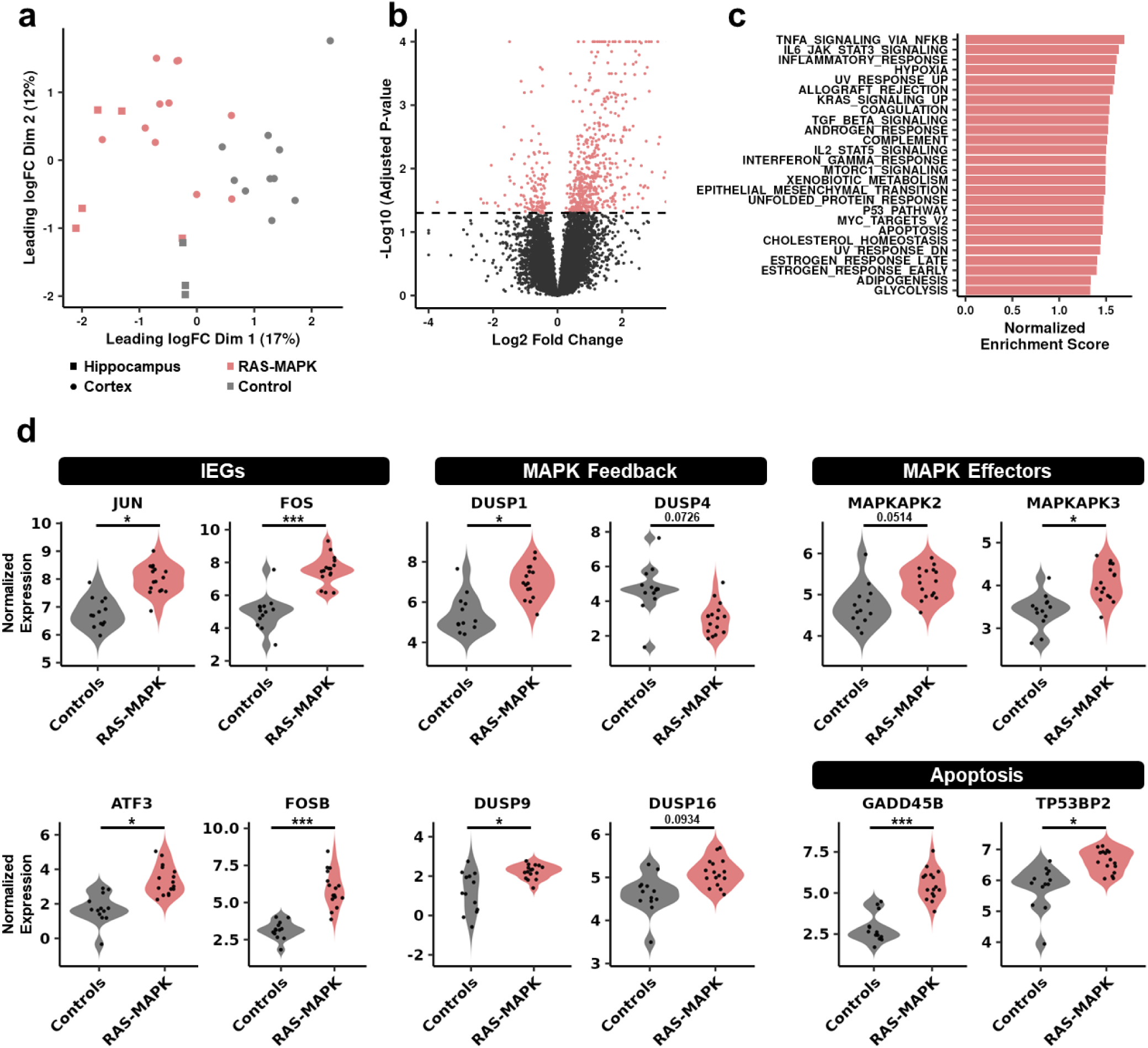
Chronic MAPK Signaling Promotes Compensatory Cellular Stress. MDS plot showing patient samples with RAS-MAPK pathway somatic findings (pink) cluster distinctly from neurotypical control samples (gray) for hippocampal (squares) and cortical (circles) tissues **(A)**. Volcano plot showing differential gene expression for patients vs. controls **(B)**. Significant pathways identified by gene set enrichment analysis included KRAS and MTORC1 signaling, as well as inflammation and apoptosis **(C)**. Violin plots showing representative findings by gene. Activation of the p38-MK2/MK3 stress response axis is evident among patient samples **(D)**. *P<0.05, ***P<0.001.

### Regional and Cellular Distribution of RAS-MAPK Pathway Somatic Variants

To understand the potential contribution of somatic RAS-MAPK pathway variants to hippocampal sclerosis and/or cortical dysplasia, we investigated the anatomical distribution of variants for each patient using the high confidence VAFs derived from deep amplicon sequencing (Fig. 4A). Interestingly, somatic variants were identified in both cortical and hippocampal tissue in four of six cases. In two cases, the causal variant was only identified in hippocampus and not in adjacent dysplastic cortex (Fig. 4B). It is difficult to entirely rule out the presence of the variant due to potential sampling bias (i.e., the sample of cells used for sequencing). However, in the majority of cases where a variant was detected in both cortical and hippocampal tissue, these observations suggest there is a common genetic contribution to both hippocampal sclerosis and cortical dysplasia.

**Figure 4:**
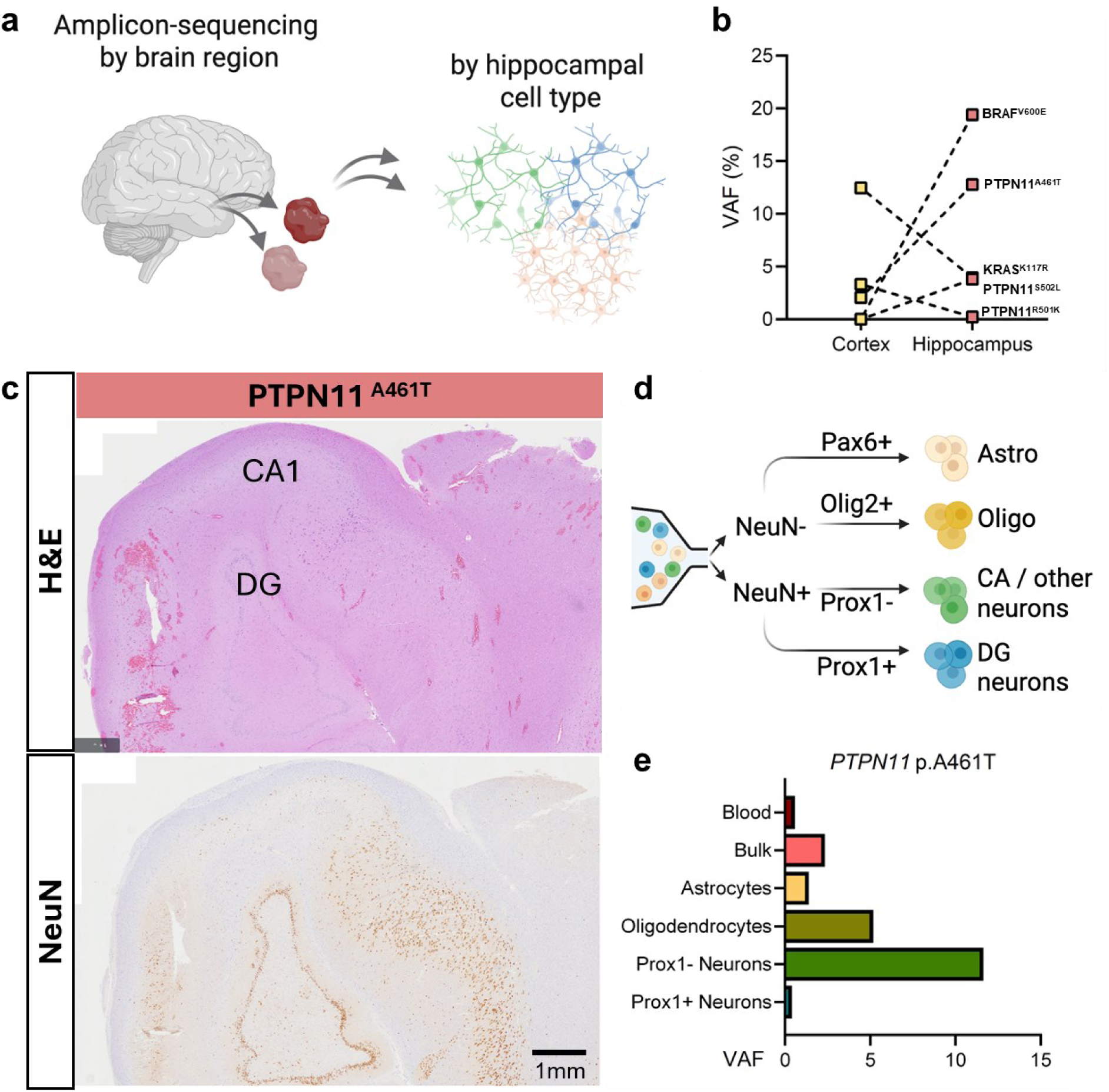
Regional and Cellular Distribution of RAS-MAPK Pathway Somatic Variants. Variant allele fractions (VAFs) from amplicon sequencing were compared across brain regions and cell types **(A)**. In four of six cases with findings, the causal somatic variant was identified in both cortex and hippocampus tissue **(B)**. One case with a PTPN11 A461T variant exhibited extensive neuronal loss in the CA1 region, as seen in H&E and NeuN stains **(C)**. Fluorescence-activated nuclei sorting was used to isolate hippocampal cell types from this tissue for amplicon sequencing **(D)**. The PTPN11 A461T variant was present in all cell types but highly enriched among Prox1-neurons **(E)**.

To identify which cell types carry RAS-MAPK pathway somatic variants, we used hippocampal tissue from a patient carrying the *PTPN11* p.A461T variant given its relatively high VAF of 12.78%. Neuropathological evaluation of this specimen showed marked loss of neurons in CA1 (Fig. 4C). We used fluorescence-activated nuclei sorting to purify hippocampal dentate granule neurons (NeuN+/Prox1+), CA1-CA3 and other neuronal subtypes (NeuN+/Prox1-), astrocytes (NeuN-/Pax6+), and oligodendrocytes (NeuN-/Olig2+) from frozen tissue (Fig. 4D). We then performed targeted amplicon sequencing to estimate VAF per cell type compared to a sample of bulk unsorted nuclei. The *PTPN11* variant was present among all cell types but highly enriched in Prox1-negative hippocampal neurons (Fig. 4E), despite this population being a minority of neurons in the sample, likely due to the marked CA1 degeneration (Ext. Data Fig. 2). This suggests that the presence of the somatic variant coincides with the affected cell types.

### *Ptpn11*^D61Y^ Mice are Susceptible to Hippocampal Degeneration Following Kainic Acid-induced Seizure

To determine whether RAS-MAPK pathway somatic variants directly contribute to hippocampal degeneration, we generated a mouse model in which Emx1-Cre drives a *Ptpn11*^D61Y^ gain-of-function variant in cortical and hippocampal excitatory neurons beginning during embryonic neurogenesis. A prior study using this model reported increases in protein tyrosine phosphatase activity and neuronal activity-induced pERK expression but did not report any spontaneous seizure activity (31). Similarly, we observed no spontaneous seizures during routine handling.

As patients in our cohort were all affected by recurrent early life seizures, we tested whether the *Ptpn11*^D61Y^ mice would be susceptible to developing hippocampal sclerosis following chemically induced seizure activity. We administered five weekly kainic acid injections starting at weaning (postnatal day (P)21). Kainic acid-induced seizures are known to elicit hippocampal degeneration in wild-type mice at high doses (32–37), so we developed a subthreshold dosing paradigm that does not induce hippocampal sclerosis in wild-type mice. An escalating dose paradigm was optimized to elicit seizure activity over the course of postnatal development, during which time sensitivity to kainic acid is known to fluctuate (38) (Fig. 5A). Both wild-type and mutant mice responded to kainic acid treatments with behavioral signs of seizure, which averaged about 1 point higher on the Hu et al. severity scale for mutant versus wild-type mice (Ext. Data Fig. 3A) (35). For both groups, seizures lasted for less than 2 hours during each session.

**Figure 5:**
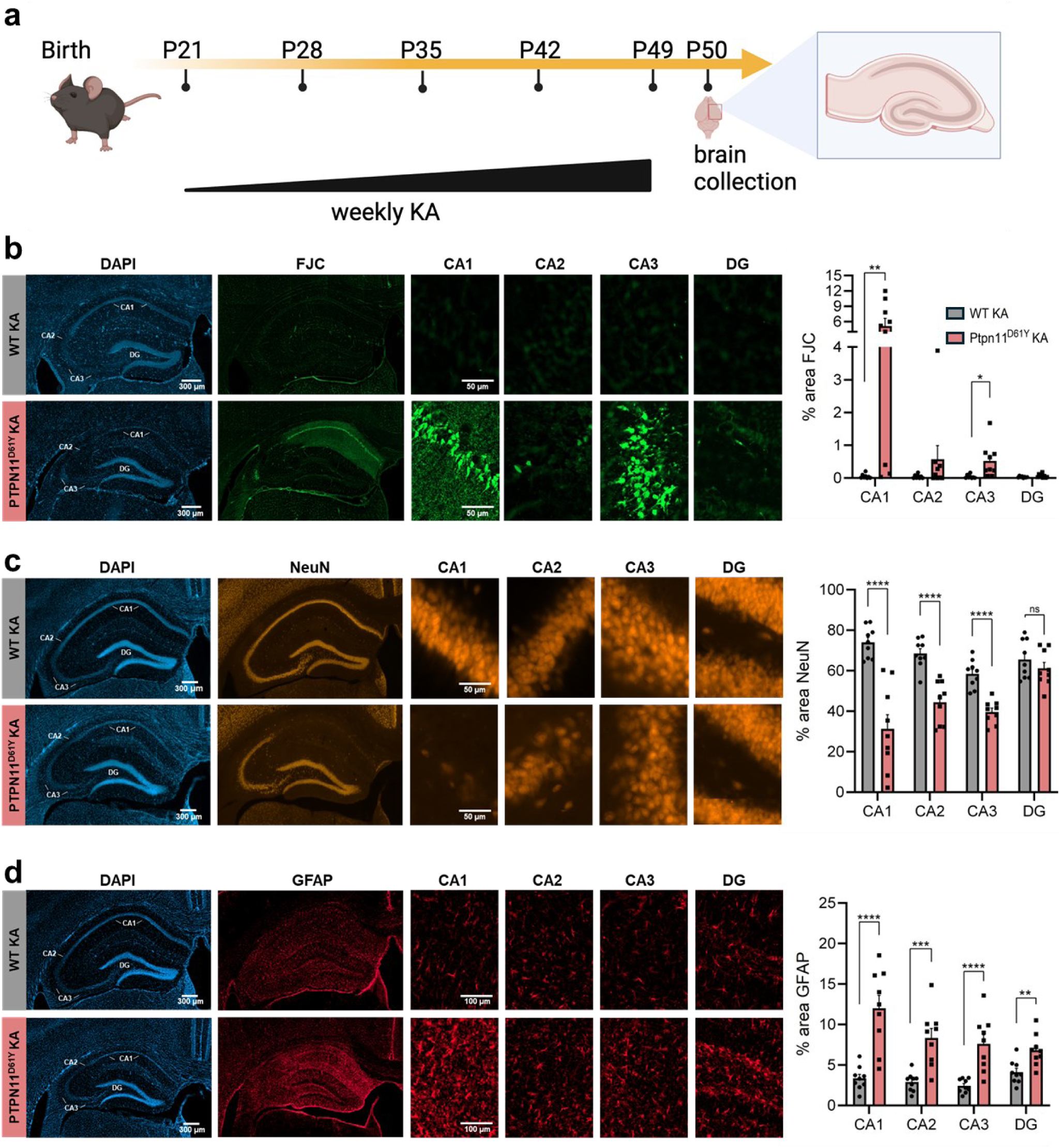
*Ptpn11*^D61Y^ mice are susceptible to hippocampal sclerosis. Low dose kainic acid (KA) treatment paradigm **(A).** Seizure induction leads to hippocampal degeneration, indicated by Fluorojade-C staining, in the CA1 and CA3 regions of mutant mice, whereas wild-type mice are unaffected **(B)**. *Ptpn11*^D61Y^ mutants have reduced NeuN staining **(C)** and gliosis (GFAP) **(D)** compared to wild-type mice. Data represent mean ± SEM (n = 8 - 9 per group; two-tailed t-tests). ns = not significant, *P<0.05, **P<0.01, ***P<0.001, ****P<0.0001.

One day after the last kainic acid injection (P50), brains were collected and examined for signs of hippocampal sclerosis. Kainic acid treatment induced extensive neuronal degeneration, indicated by Fluorojade-C staining, in the CA1 and CA3 regions of the hippocampus of *Ptpn11*^D61Y^ mice, whereas wild-type mice were unaffected (Fig. 5B). Moreover, *Ptpn11*^D61Y^ mice had significant reductions in NeuN-positive neurons in CA1, CA2, and CA3 (Fig. 5C), as well as marked signs of gliosis as indicated by GFAP staining (Fig. 5D). There were no significant correlations between seizure severity score and neuronal degeneration (Ext. Data Fig. 3B-D), indicating individual differences in seizures were not driving the increased damage seen in mutant mice. However, the extent of neuronal damage was strongly correlated to the extent of gliosis observed in the CA1, CA2, and CA3 regions (Ext. Data Fig. 4). Of note, this neuronal loss was specific for the hippocampus. No signs of neuronal degeneration were observed in the cortex of mice belonging to either group, even though Emx1-Cre drives expression of the *Ptpn11*^D61Y^ variant in cortical excitatory neurons in addition to hippocampal neurons (Ext. Data Fig. 5).

Though past studies reported no gross anatomical changes in the hippocampus of this mouse model (31), we assessed baseline hippocampal integrity by staining brains with Fluorojade-C, NeuN, and GFAP after 5-weeks of saline treatment (Ext. Data Fig. 6A). Mice showed no signs of degeneration or gliosis, demonstrating that the genetic variant alone is not sufficient to cause these changes (Ext. Data Fig. 6B-D). Taken together, these results suggest that *Ptpn11* gain-of-function sensitizes mice to developing hippocampal sclerosis in the presence of seizure activity.

**Figure 6:**
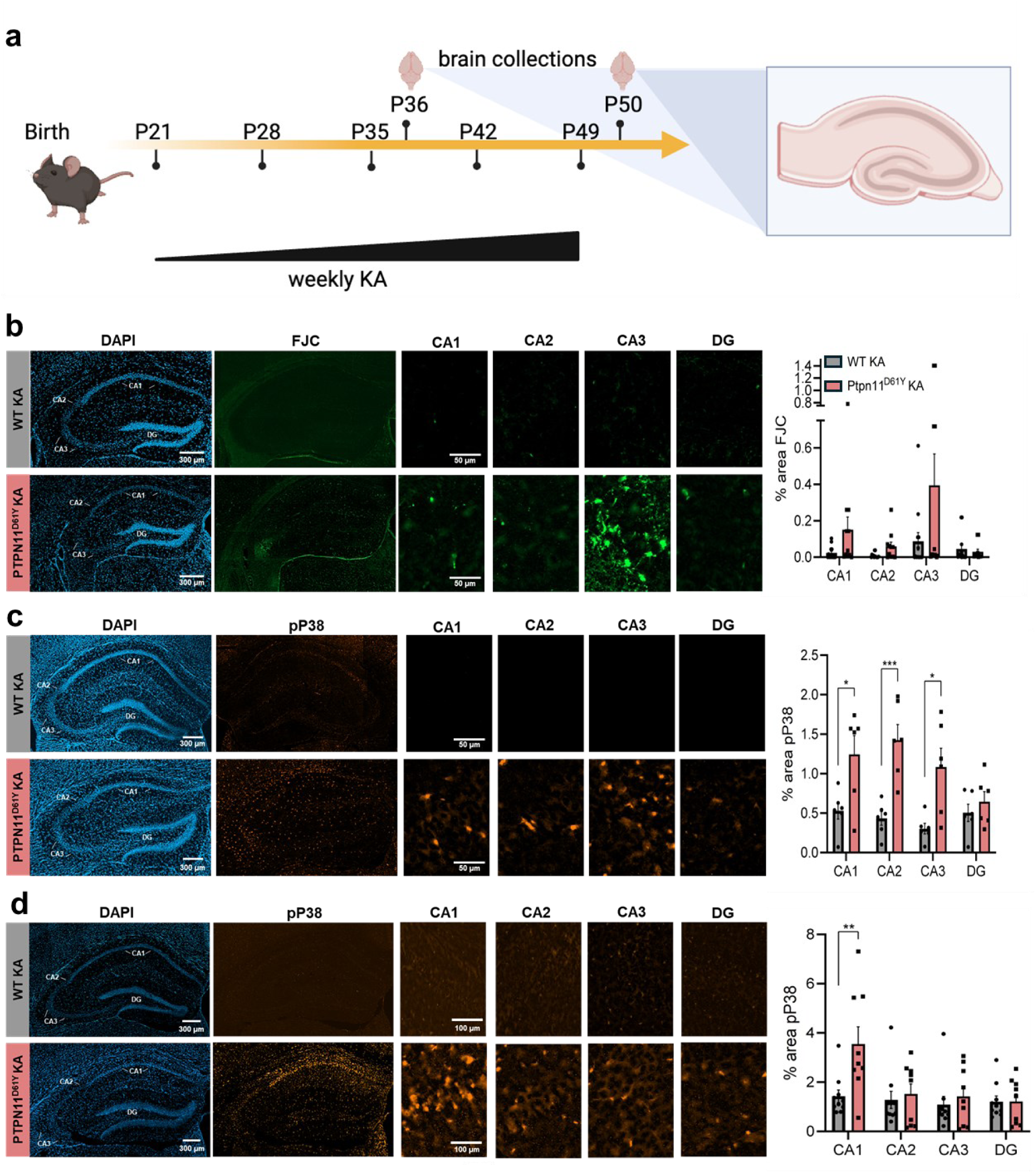
Accumulated Cellular Stress is Associated with Susceptibility in *Ptpn11*^D61Y^ Mice. A separate cohort of mice was collected at P36, following the first behavioral seizure, to assess the time course of degeneration **(A)**. Minimal Fluorojade-C staining was evident at this timepoint **(B)**. Phospho-p38 staining was evident at P36 **(C)**, suggesting cellular stress was already present, and it increases in CA1 by P50 **(D)**. Data represent mean ± SEM (n = 6 - 13 per group; two-tailed t-tests). *P<0.05, **P<0.01. KA = kainic acid

### Accumulated Cellular Stress is Associated with Susceptibility in *Ptpn11*^D61Y^ Mice

To better understand the mechanism underlying this susceptibility, we asked when neuronal damage occurs in *Ptpn11*^D61Y^ mice, which would inform whether it represents an acute excitotoxic response or a gradual cumulative effect. We collected brains from a separate cohort of mice one day following the third kainic acid injection (P36), which also corresponds to the first behavioral seizure in this model (Fig. 6A). *Ptpn11*^D61Y^ mice showed minimal Fluorojade-C staining which was not significantly different from control levels, although a few mutant mice had noticeable Fluorojade-C positivity in CA3, suggesting that they are already more vulnerable to seizure-induced hippocampal damage at this early timepoint, but that damage accumulates over time (Fig. 6B).

Given the gene expression changes noted in patient tissues, which implicated cellular stress through the p38 MAPK pathway, we asked whether phospho-p38 expression was increased in *Ptpn11*^D61Y^ mice. After the third kainic acid injection, we observed broadly increased phospho-p38 in mutant mice (Fig. 6C). At the height of neuronal degeneration (5 weeks of kainic acid injections), we observed strongly elevated phospho-p38 staining in CA1 of the hippocampus of mutant mice relative to controls, although other regions showed similar levels as controls (Fig. 6D). This suggests p38-mediated cellular stress precedes neuronal cell death. At baseline, saline-treated mice of both genotypes exhibited no phospho-p38 staining, again demonstrating an interaction of seizures with genotype (Ext. Data Fig. 6E). Taken together, these findings suggest that *Ptpn11*^D61Y^ mutants sustain elevated p38 MAPK signaling which leads to an accumulation of hippocampus-specific cellular stress and damage over repeated seizure episodes, which increases susceptibility to developing hippocampal sclerosis.

## Discussion

Hippocampal sclerosis has traditionally been conceptualized as an acquired pathology, particularly in FCD IIIa where seizures originating from cortical dysplasia may cause hippocampal damage (39, 40). Cortical dysplasia can be observed from infancy, whereas hippocampal sclerosis is rarely observed prior to 2 years of age, suggesting it develops over time (11). In addition, a history of febrile seizures or febrile status epilepticus as an initial precipitating injury is a strong predictor of hippocampal sclerosis (41). Despite this evidence, earlier studies have hinted at a possible genetic component. For example, children with childhood convulsions who progress to develop hippocampal sclerosis have subtle asymmetries already evident by brain imaging before the age of two (11). In addition, rare familial occurrence of hippocampal sclerosis has been reported (42). Our results clarify this relationship by demonstrating that both precipitating events and genetic components are implicated in the development of hippocampal sclerosis, such that somatic variants activating the RAS-MAPK pathway confer susceptibility in the presence of seizure activity.

Recent studies identified post-zygotic somatic variation of the RAS-MAPK pathway in approximately 10% of patients with MTLE, most of whom were adults (9). Hippocampal sclerosis is a common histopathological finding in drug-resistant temporal lobe epilepsy both in children and adults, but there are some important distinctions (43). In the pediatric setting, hippocampal sclerosis frequently coincides with cortical dysplasia and early-onset epilepsy as FCD IIIa in as many as 79% of cases (10, 44–49). This contrasts the adult population, in which two-thirds of patients with temporal lobe epilepsy have standalone hippocampal sclerosis without cortical dysplasia and a later age of seizure onset (10, 50). Given a common link to somatic variants in both presentations, it is possible that these two entities are part of a spectrum and the differences in brain regions implicated (i.e., primarily hippocampus in MTLE and both hippocampus/cortex in FCD IIIa) could reflect a difference in the developmental origin of these somatic variants. Variants arising earlier in brain development in a cellular lineage common to the temporal cortex and hippocampus may lead to FCD IIIa, whereas variants originating later in development in a lineage exclusive to the hippocampus may lead to adult-onset MTLE. Indeed, Khoshkhoo et al. observed RAS-MAPK pathway somatic variants restricted to hippocampus in their patient cohort (9, 51). In our cohort, variants were found in both cortex and hippocampus. Our data together with the prior literature suggest that somatic variants activating the RAS-MAPK pathway are implicated in a spectrum of disorders encompassing both adult mesial temporal lobe epilepsy and FCD IIIa.

MAPK signaling represents a highly complex, branched network with extensive regulation, feedback, and cell type-specific effects. Although RAS-MAPK variants are known to alter the canonical ERK pathway, our gene expression analysis of patient tissues also showed strong evidence of p38 MAPK activity, which is known to reflect inflammation, cellular stress, and neuronal vulnerability in the hippocampus. In animal models, neuronal loss typically correlates with seizure severity, with stress-activated pathways such as p38 MAPK acting as downstream effectors of excitotoxic damage (52–55). However, *Ptpn11*^D61Y^ mice exhibited only modestly increased seizure severity following kainic acid, yet they showed disproportionately elevated p38 activation and hippocampal neuronal loss. In neuronal cell culture models, sustained ERK activity has been shown to directly induce p38 signaling and cell death (56). GADD45β, a key mediator of ERK-p38 crosstalk, was strongly upregulated in our patient brain tissues, suggesting a direct link between RAS-MAPK variants and increased p38 signaling (57). These observations raise the possibility that RAS-MAPK variants may lower the threshold for seizure-induced injury by engaging p38-dependent stress pathways. This model challenges the conventional view that neuronal degeneration is solely determined by seizure burden and instead suggests that genetic context shapes the brain’s vulnerability to excitotoxic damage.

Importantly, our results raise the possibility of precision medicine targeting the MAPK pathway in drug-resistant epilepsy. Treatment options for FCD IIIa include standard antiseizure medications and surgical resection. However, many patients have seizures that are resistant to medication, and not all patients are candidates for surgery. In some cases, resection may lead to negative consequences including memory or language deficits. In other cases, seizures may recur after surgery. Identifying a molecular cause of FCD IIIa provides new opportunities to pursue targeted therapeutics. For example, MAPK pathway inhibitors could potentially be repurposed for epilepsy or considered as a preventative therapy for patients presenting with febrile status epilepticus, a known risk factor for hippocampal sclerosis. Taken together, these results provide a plausible mechanism for both the prevention of hippocampal sclerosis in people with genetic risk as well as shifting the treatment paradigm away from surgery and towards pathway-specific interventions.

## Methods

### Study Participants

Neurosurgical patients treated for drug-resistant epilepsy at Nationwide Children’s Hospital were enrolled via written consent on an Institutional Review Board (IRB)-approved research protocol. Prior to surgery, each patient was evaluated for neuroimaging and neurophysiology by a multidisciplinary pediatric clinical care team. Blood and brain tissue samples within the epileptogenic zone were collected at the time of surgical resection and defined by anatomic location, with each block of tissue divided for clinical neuropathologic evaluation and snap frozen for genomic analysis. In some cases, formalin-fixed tissue was also collected for genomic analysis. The neuropathology review of adjacent formalin-fixed paraffin-embedded tissue sections was performed by board-certified neuropathologists according to International League Against Epilepsy (ILAE) guidelines. Neuropathological diagnoses were confirmed by immunohistochemical markers when indicated for clinical evaluation. Resected hippocampal and cortical tissues, as available, were sequenced from patients with a diagnosis of mesial temporal sclerosis or hippocampal sclerosis indicated by neuroimaging and/or neuropathology. Resected hippocampal tissue from patients without any evidence of sclerosis and postmortem donor tissue from NIH NeuroBioBank served as controls.

### Exome Sequencing and Analysis

Exome sequencing libraries were prepared from 500 ng of DNA isolated from blood and brain tissue using the NEBNext Ultra II FS DNA Library Prep Kit (New England Biolabs). Libraries were pooled for hybrid exome capture using the IDT xGen Exome Research Panel v1.0 kit enhanced with the xGenCNV Backbone Panel-Tech Access (Integrated DNA Technologies). Final libraries were sequenced on an Illumina HiSeq 4000 or NovaSeq 6000 to generate paired-end 151-bp reads. Reads were mapped to human reference genome build GRCh38 using bwa v0.7.15 and secondary analysis performed under the Churchill framework (58). Sequence alignments were refined according to community-accepted guidelines for best practices (https://www.broadinstitute.org/gatk/guide/best-practices).

Detailed quality control metrics were computed for quality control purposes. For each base pair, the total sequence generated, mapped/unmapped, soft-clipped, and marked duplicate were computed from the BAM files using Picard (https://broadinstitute.github.io/picard/) v2.17.11. Depth and breadth of exome sequencing coverage across target regions were computed from the aligned BAM8 files by MosDepth using target region BED files provided by the kit manufacturer (IDT) (59). The biological sex of each sample, determined by computing the ratio of reads aligned to chrX versus chrY (>75:1 considered female, <20:1 considered male, ratios between called ambiguous) was compared to the sex at birth from patient medical record. Sample contamination was estimated using VerifyBamId v1.1.3 (60).

Somatic variants in tissue samples were called using MuTect-2 using the blood sample as matched normal comparator (61). Somatic variants were filtered based on the following features: MuTect-2 filter = “PASS”, GATK quality score (≥30), depth of sequencing (≥eight total reads), absence from blood comparator, alternate allele reads (≥4), and variant allele frequency (≥1%). Variants that passed these initial filters were visually reviewed in Integrated Genomics Viewer (IGV) to remove likely artifactual calls arising from sequencing error, strand bias, or local read misalignment. Variants passing visual review were annotated using snpEff, filtered for rare (MAF < 0.1% in gnomAD v. 4.1 and/or All Of Us databases) non-synonymous, truncating, and splice-site variants. Somatic copy number variation was called using VarScan2 v2.4.3 (parameters: --min-coverage 50 --min-segment-size 10 --max-segment-size 100) using exome sequencing data from blood as a comparator for each brain tissue sample (62). Copy number segments with num.mark ≥30 and adjusted log2 value either <−0.25 or >0.25 were annotated with RefSeq gene exons to aid interpretation.

### Amplicon Sequencing

Somatic variants were validated by amplicon sequencing, as described previously (4, 7, 63). Human reference DNA GM24143 or HG04217 (Coriell Institute) was used as a negative control. Primer sequences for targeted amplification are provided in Supplementary Table 5. SAMtools mpileup and mpileup2cns VarScan (version 2.3.4) commands were used to generate read counts for variants of interest from raw, de-duplicated BAM files.

### Protein Structure Modeling and Identification of Tertiary-Level Interactions

Protein structure modeling of the PTPN11 protein tyrosine phosphatase domain and KRAS variants was performed using ColabFold v1.5.5: AlphaFold2 using MMseq2 using no supplied template and under default settings (64). Structural models were examined using the Cystallographic Object-Oriented Toolkit for Windows (65) to identify predicted interactions formed by residues of interest using the Environmental Distances tool with a cut-off of 4.0Å.

### RNA Sequencing and Differential Expression Analysis

Total RNA was extracted from frozen brain tissue using the Qiagen AllPrep kit. Ribosomal RNA depletion and library construction for RNA sequencing was performed with NEBNext rRNA Depletion Kit and the NEBNext Ultra II Directional RNA Library Prep Kit. Two patient samples were prepped using the Watchmaker RNA Library Prep Kit for Illumina. No batch effect related to library kit was observed. Sequencing was performed on an Illumina HiSeq 4000 or NovaSeq 6000 according to manufacturer protocols. Reads were mapped and quantified to GENCODE Human Release 28 transcriptome using salmon v1.9.0, and bootstrapping = 100. Results of Salmon quantification were consolidated from transcript to gene level and were analyzed using differential expression for repeated measures (dream) in R, which utilizes a linear mixed model for datasets with nested samples per individual (66). Dream was run using a mixed model including the disease state of samples (case or control) and patient ID: ∼disease_state + (1|patient_id). Differentially expressed genes (DEGs) were identified as having a Benjamini-Hochberg FDR ≤ 0.05. Volcano and violin plots were generated in R using the ggplot2 package.

Ensembl gene ids (ENSG) of DEGs were submitted to The Database for Annotation, Visualization, and Integrated Discovery (DAVID), using the full list of ENSG from Dream as the background. Enriched GO terms were identified from GOTERM BP DIRECT using a Benjamini-Hochberg FDR of ≤ 0.05. The fgsea R package was used to perform gene set enrichment analysis against the MsigDB human Hallmark gene set, v2024.1 with 200,000 permutations, using signed adjusted p-values (Log2 FC * - Log10(adj.P.Val)). Significantly enriched gene sets were identified by a Benjamani-Hochberg FDR of ≤ 0.05 and plots were generated using R and the ggplot2 package.

### Fluorescence-Activated Nuclei Sorting

Nuclei were isolated from fresh frozen tissue samples using the Chromium Nuclei Isolation Kit (10X Genomics). Nuclei were labeled using the following antibodies: NeuN (Millipore, #MAB377X), PROX1 (Novus Biologicals, #NBP1-30045AF647), OLIG2 (Novus Biologicals, # NBP2-89201AF594), and PAX6 (Novus Biologicals, #NBP2-34705AF647) at 1:100 followed by incubation for 1 hour at 4°C with agitation. All nuclei were then stained with DAPI (ThermoFisher Scientific #EN62248). Cell types of interest (NeuN+/PROX1-: neurons, NeuN+/PROX1+: hippocampal dentate granule neurons, NeuN-/OLIG2+: oligodendrocytes, NeuN-/PAX6+: astrocytes) were gated and sorted using a Bigfoot Cell Sorter running Sasquatch 1.19.3 software (ThermoFisher Scientific). DNA was extracted from sorted nuclei using QIAamp DNA Micro Kit (Qiagen). The extracted DNA was used for high-depth targeted sequencing as described above.

### Immunofluorescence – Patient Tissue

FFPE tissue was deparaffinized in xylenes and rehydrated in a gradient of ethanol washes. Slides were then washed (PBS with 0.025% Triton X-100) and blocked in PBS with 10% donkey serum and 0.3% Triton X-100 for 1 hour. Sections were then incubated with primary antibody [(Phospho-p44/42 MAPK (Erk1/2) (Thr202/Tyr204), Cell Signaling 9101S; 1:500)] diluted in PBS with 10% donkey serum and 0.1% Triton X-100 overnight at 4°C. The next day, slides were washed and then incubated with secondary antibody [(Donkey anti-Rat IgG (H+L) Highly Cross-Adsorbed Secondary Antibody, Alexa Fluor™ Plus 647; 1:1,000)] and DAPI diluted in PBS with 10% donkey serum and 0.1% Triton X-100 for 2 hours at room temperature. Slides were then washed and mounted with ProLong Diamond Anti-Fade mounting medium (ThermoFisher). Slides were imaged with an Axio Imager M2 (Zeiss).

### Experimental animals

Experimental mice were of C57BL6 background and were group-housed on a 12-h light/dark cycle (lights on at 6:00 a.m.) with access to standard chow and water *ad libitum*. All experiments were approved and performed in compliance with the animal care guidelines issued by the National Institutes of Health and the institutional animal care and use committee of Nationwide Children’s Hospital. The mouse strain used for this research project, B6.129S6-*Ptpn11^tm6Bgn^*/Mmjax, RRID:MMRRC_032103-JAX, was obtained from the Mutant Mouse Resource and Research Center (MMRRC) at The Jackson Laboratory, an NIH-funded strain repository, and was donated to the MMRRC by Benjamin G. Neel, M.D., Ph.D., Ontario Cancer Institute (67). Heterozygous *Ptpn11*^D61Y^ floxed mice (Jackson Labs, #032103) were crossed with homozygous *Emx1*-Cre (Jackson Labs, #005628) to generate litters containing on average 50% WT *Ptpn11* mice and 50% heterozygous *Ptpn11*^D61Y^ gain of function mice. Mice were genotyped for the *Ptpn11* variant by PCR using primers ACGCCTTCTCTCAATGGACT and GCAGACCTGGTCACGAAGAT. PCR reactions were purified using the Qiaquick PCR Purification Kit (Qiagen) followed by Sanger sequencing (Eurofins).

### Seizure Induction and Behavioral Scoring

Kainic acid (Cayman Chemical) was resuspended in a 1x phosphate buffered saline (PBS) solution (Fisher Scientific). Mice were administered increasing doses of kainic acid or an equivalent volume of PBS (vehicle control) via intraperitoneal (i.p.) injection once every week for 3 or 5 weeks from P21 – P49. Mice were observed continuously for at least 1 hour and assigned a seizure severity score based on the animal’s behavior as follows: grade 0: no response; grade I: staring, limbs pawing; grade II: staring, pawing, nodding; grade III: staring, nodding, pawing, rearing, falling over, jumping; grade IV: status epilepticus and death (35). At P36 or P50, mice were anesthetized using a cocktail of ketamine (300 mg/kg) and xylazine (15 mg/kg) and while under a deep plane of anesthesia mice were transcardially perfused with ice cold PBS followed by 4% paraformaldehyde. Brains were collected and post-fixed in 4% paraformaldehyde for at least 24 hours at 4°C and then transferred to 30% sucrose, 0.02% sodium azide and stored at 4°C.

### Immunofluorescence – Mouse Tissue

Coronal sections at 30 µm thickness were collected via a freezing microtome (Leica Cat# SM2010R) and stored in 1x PBS with 0.02% sodium azide. For Fluoro-Jade C staining, sections were mounted on 1% gelatin coated slides using 1X EZ-Float®Tissue Mounting Reagent (Millipore Sigma) and dried overnight on a slide warmer at 50°C. Slides were stained using the Fluoro-Jade C Staining Kit with DAPI Counter Stain (Histo-Chem, Inc) according to the manufacturer’s protocol and imaged with an Axio Imager M2 (Zeiss). For immunofluorescence, free floating sections were washed (PBS with 0.025% Triton X-100) and blocked in PBS with 10% donkey serum and 0.3% Triton X-100 for 1 hour. Sections were then incubated with primary antibodies for NeuN (Alexa Fluor™ 555 conjugate, Millipore Sigma, MAB377A5; 1:500), GFAP (Cell Signaling, 80788; 1:500), and Phospho-p38 MAPK [(Thr180/Tyr182) (D3F9) Cell Signaling, 4511; 1:1000] diluted in PBS with 10% donkey serum and 0.1% Triton X-100 overnight at 4°C. The next day, sections were washed and then incubated with secondary antibody [(Donkey anti-Rabbit IgG (H+L) Highly Cross-Adsorbed Secondary Antibody, Alexa Fluor™ Plus 647 or 555; 1:1,000)] and DAPI diluted in PBS with 10% donkey serum and 0.1% Triton X-100 for 2 hours at room temperature. Sections were then washed and mounted using 1X EZ-Float®Tissue Mounting Reagent (Millipore Sigma) and ProLong Diamond Anti-Fade mounting medium (ThermoFisher).

### Image Analysis

Slides were imaged with an Axio Imager M2 (Zeiss) using consistent settings within experiments. To quantitate the degree of cellular loss and reactive gliosis, CZI files were imported into ImageJ Fiji (version 1.54p), and morphological landmarks were used to manually annotate regions of interest (ROIs) encapsulating CA1, CA2, CA3, and the dentate gyrus. Rolling ball background subtraction (50 pixel radius) was applied to Fluor-Jade C, GFAP, and phospho-P38 and median blur (2 pixel radius) were applied to the fluorescence channels containing the markers of interest as standard pre-processing steps prior to segmentation with the Otsu (NeuN, GFAP), Yen (Fluoro-Jade C), or Triangle (phosho-P38) thresholding algorithms in ImageJ. The “Analyze Particles” function was used to identify the percent area of segmented signal within each ROI. Phospho-P38 measurements for % area was taken by segmentation of p38 signal with a minimum size of 10 um and 0.15-1 circularity to identify cells/nuclei with significant upregulation of phosphor-P38.

### Statistical Analysis

Statistical tests are indicated in the main text and figure legends. P≤0.05 was considered statistically significant.

## Acknowledgments

We would like to thank the patients and families for contributing their specimens to advance our understanding of the genetic bases of epilepsy. This work was supported in part by NIH-NINDS (R01NS129784) to TAB. The content is solely the responsibility of the authors and does not necessarily represent the official views of the NIH. Some figures were created in BioRender.

## Author Contributions

LMW and TAB conceptualized the study. LMW, AH, KS, SS, AR, AT, JA, CHS, EARG, SR, ARZ, KEM, and DCK performed experiments and/or analyzed data. TAB and ERM contributed funding and resources. AD and MC contributed to patient consent and sample procurement. JL, JP, AS, DRB, DLT, CRP, APO contributed to clinical characterization of the study cohort. LMW and TAB wrote the original draft. All authors participated in review and editing of the manuscript.

## Competing Interests Statement

The authors report no conflicts of interest.

---

**Ext. Data Figure 1:**
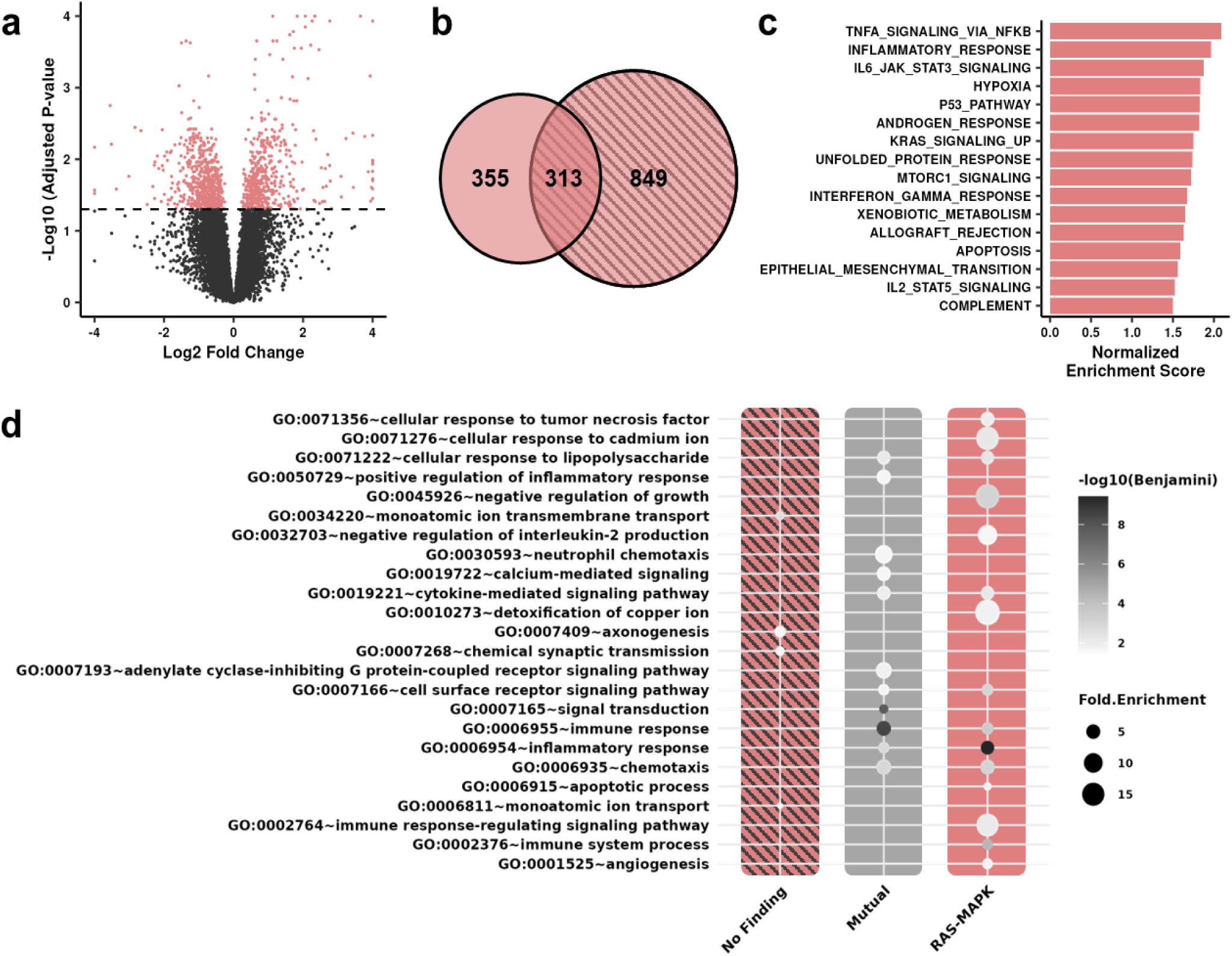
Gene expression comparison of cases with and without genetic findings. Volcano plot showing differential gene expression between cases with no findings vs. neurotypical controls **(A)**. Venn diagram of overlapping DEGs resulting from comparisons of patients with RAS-MAPK variants vs. controls (left) and patients without findings vs. controls (right) **(B)**. Gene set enrichment analysis of DEGs identified in cases without genetic findings **(C)**. Gene ontology terms identified in genes shared among comparisons or exclusive to either patient subset **(D)**.

**Ext. Data Figure 2:**
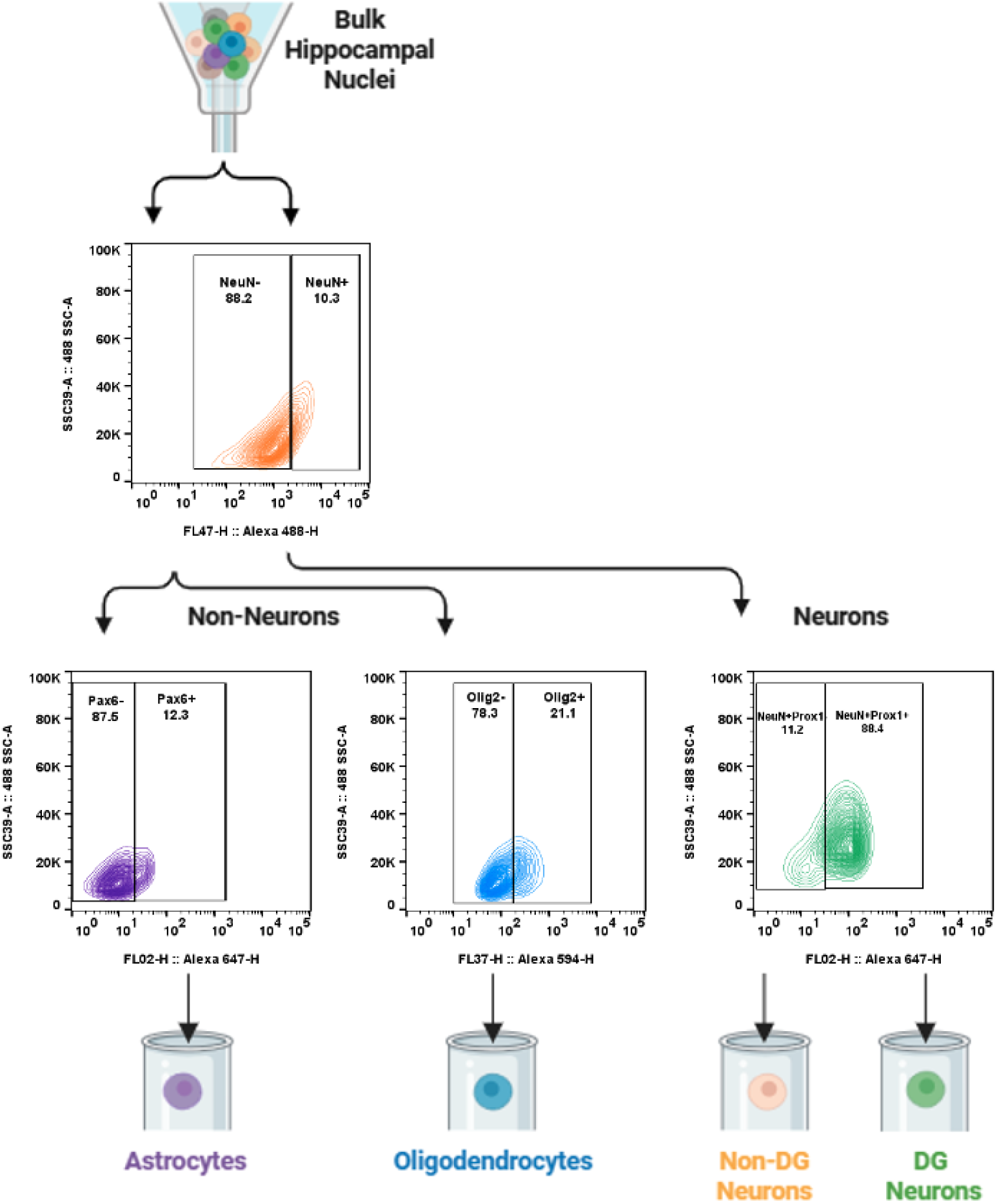
Fluorescence-activated nuclei sorting. Sorting scheme to enrich astrocytes, oligodendrocytes, Prox1-negative neurons and Prox1-positive neurons for amplicon sequencing.

**Ext. Data Fig. 3:**
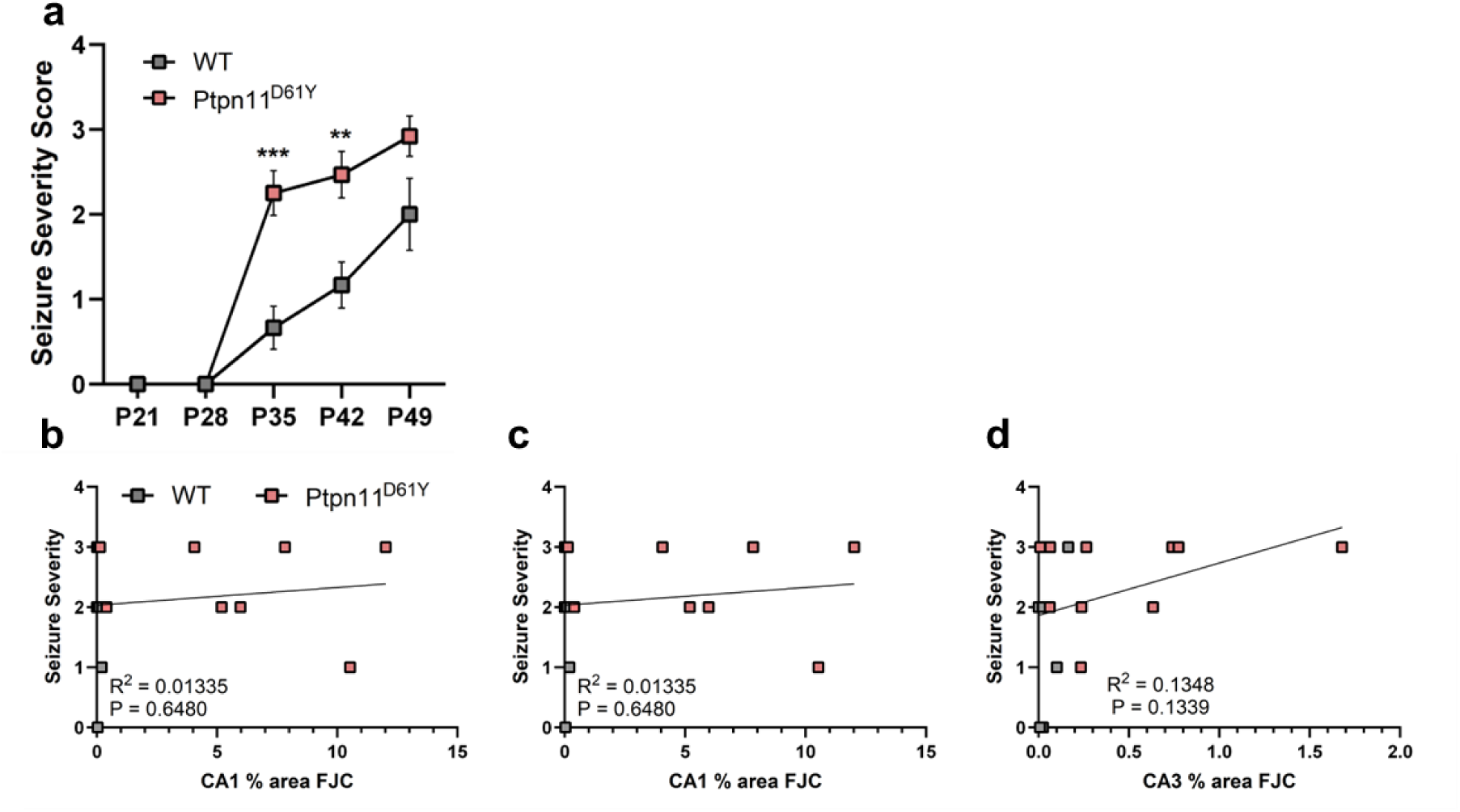
Seizure severity does not drive neuronal damage. Behavioral scoring of seizure severity after each kainic acid injection. Mutant mice scored about 1-point higher than controls. Note: among *Ptpn11*^D61Y^ mice, 1 died at P35, 2 at P42, and 3 at P49. For wild-type mice, 2 died at P49. Data represent mean ± SEM (n = 12 - 16 per group; Mann-Whitney test). **P<0.01, *** P<0.001 **(A)**. Seizure severity did not correlate with Fluorojade-C staining, suggesting individual differences in seizure severity do not drive the degenerative effect observed in mutants **(B-D)**.

**Ext. Data Fig. 4:**
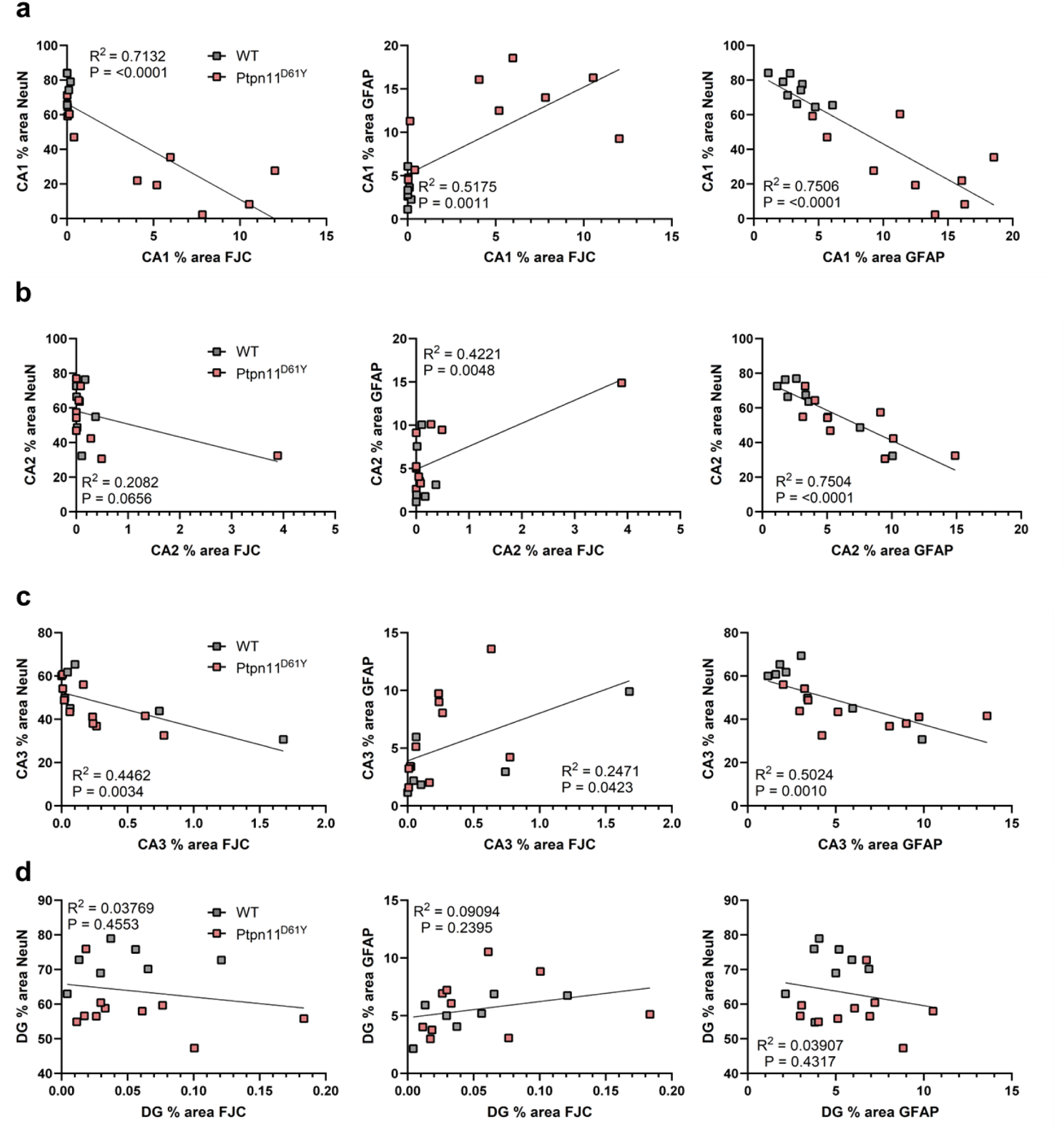
Correlation among hippocampal outcome measures. The extent of neuronal loss and gliosis tend to correlate across affected brain regions, including CA1, CA2, and CA3 **(A-C)**, except for the dentate gyrus which was unaffected **(D)**.

**Ext. Data Fig. 5:**
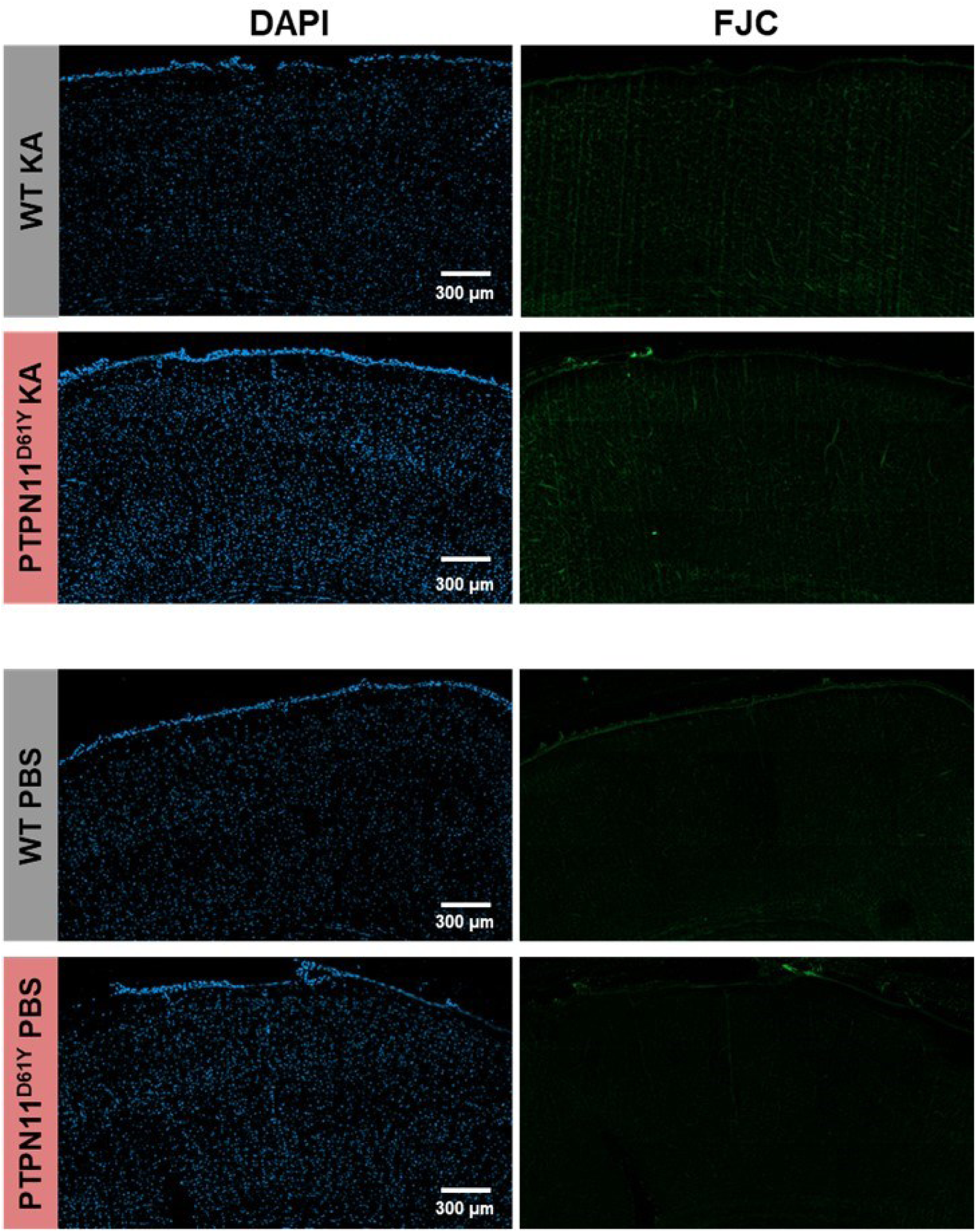
Neuronal degeneration is specific for the hippocampus. No Fluorojade-C staining was observed outside the hippcampus. Primary somatosensory cortex is shown for both genotypes following the kainic acid (KA) treatment paradigm (top) and saline treatment paradigm (bottom).

**Ext. Data Fig. 6:**
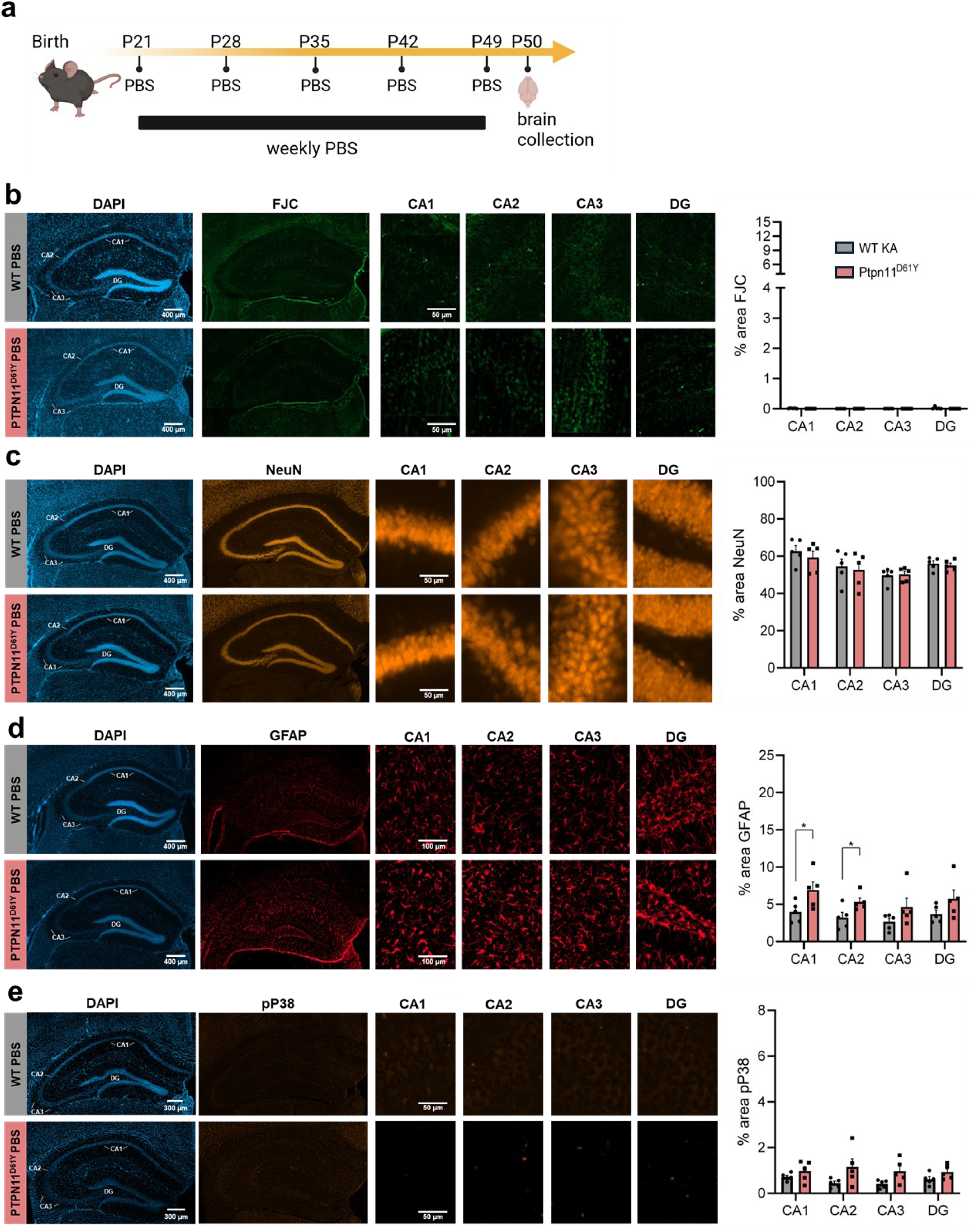
Baseline hippocampal integrity is similar in WT and PTPN11 mutant mice. Mice were treated with five weekly i.p. injections of phosphate-buffered saline (PBS) from P21 to P49 and brains were examined one day later at P50 **(A)**. There was no sign of neuronal degeneration in the hippocampus of either genotype based on Fluorojade-C staining **(B)** or NeuN staining **(C)**. *Ptpn11* ^D61Y^ mutants had slightly more GFAP staining (∼2% more) in CA1 and CA2 regions compared to wild-type mice **(D)**. Phosho-p38 expression was similarly absent among groups **(E)**. Data represent mean ± SEM (n = 5 per group; two-tailed t-tests). *P<0.05.

## Supplementary Text: Structural modeling of PTPN11 variants

For wildtype PTPN11, we modeled the protein-tyrosine phosphatase domain and found that A461 colocalizes with R501, and S502 within the tertiary in proximity to the catalytic pocket housing the catalytic cysteine, C459. Interestingly, the backbone of A461 is predicted to form a hydrogen-bond (3.4Å) with the sidechain of R501. We then modeled the PTPN11 protein-tyrosine phosphatase domain with each of the validated patient somatic variants and each predicted structural model was found to closely align with their respective wildtype model: PTPN11^A461T^ (0.054), PTPN11^R501K^ (0.042), PTPN11^S502L^ (0.126). To better highlight the changes in tertiary interactions, interactions formed by the backbones of these residues which are predicted to be retained were excluded from the analysis. In the wildtype PTPN11 protein-tyrosine phosphatase domain, the backbone of A461 is predicted to interact with the backbone of I463 (3.8Å). Substitution of this hydrophobic alanine sidechain with polar threonine (A461T) is predicted to disrupt this interaction in favor for a shorter interaction between the sidechain of T461 and the backbone of I463 (3.4Å). Additionally, the hydroxyl group is predicted to form interactions with the backbone of G462 (3.1Å) and sidechain of Q506 (3.7Å), predictively introducing new interactions between residues forming the catalytic pocket. The sidechain of R501 within the wildtype PTPN11 protein-tyrosine phosphatase domain is predicted to form numerous interactions with neighboring residues, including the backbones of A461 (3.4Å) and G462 (3.0Å), as well as two interactions with the sidechain of N306 (3.1Å and 3.2Å) and sidechain of N308 (2.7Å). Resulting from the R501K mutation, multiple interactions are predicted to be abolished, most notable to A461 and G462 forming the architecture of catalytic pocket. In addition, K501 is also predicted to retain only one interaction with the sidechain of N306 (3.4Å) and its interaction with N308 (3.4Å), although the greater distance of these interactions predicts they will be weaker. The sidechain of S502 within the wildtype PTPN11 protein-tyrosine phosphatase domain is predicted to form a single interaction with the backbone of L283 (3.6Å). S502L is predicted to break this interaction, as well as alter the position of R501 but this is not predicted to significantly alter interactions made by the R501 sidechain.

## Notes

### Competing Interest Statement

The authors have declared no competing interest.

### Summary of Updates

The journal that was listed for submission was deleted at the top of page 1.

## References

1. Barba C, Specchio N, Guerrini R, Tassi L, De Masi S, Cardinale F, et al. Increasing volume and complexity of pediatric epilepsy surgery with stable seizure outcome between 2008 and 2014: A nationwide multicenter study. Epilepsy Behav. 2017;75:151–7.

2. Blumcke I, Spreafico R. An international consensus classification for focal cortical dysplasias. Lancet Neurol. 2011;10(1):26–7.

3. Najm I, Lal D, Alonso Vanegas M, Cendes F, Lopes-Cendes I, Palmini A, et al. The ILAE consensus classification of focal cortical dysplasia: An update proposed by an ad hoc task force of the ILAE diagnostic methods commission. Epilepsia. 2022;63(8):1899–919.

4. Bedrosian TA, Miller KE, Grischow OE, Schieffer KM, LaHaye S, Yoon H, et al. Detection of brain somatic variation in epilepsy-associated developmental lesions. Epilepsia. 2022;63(8):1981–97.

5. Bonduelle T, Hartlieb T, Baldassari S, Sim NS, Kim SH, Kang HC, et al. Frequent SLC35A2 brain mosaicism in mild malformation of cortical development with oligodendroglial hyperplasia in epilepsy (MOGHE). Acta Neuropathol Commun. 2021;9(1):3.

6. Lopez-Rivera JA, Leu C, Macnee M, Khoury J, Hoffmann L, Coras R, et al. The genomic landscape across 474 surgically accessible epileptogenic human brain lesions. Brain. 2023;146(4):1342–56.

7. Miller KE, Koboldt DC, Schieffer KM, Bedrosian TA, Crist E, Sheline A, et al. Somatic SLC35A2 mosaicism correlates with clinical findings in epilepsy brain tissue. Neurol Genet. 2020;6(4):e460.

8. Winawer MR, Griffin NG, Samanamud J, Baugh EH, Rathakrishnan D, Ramalingam S, et al. Somatic SLC35A2 variants in the brain are associated with intractable neocortical epilepsy. Ann Neurol. 2018;83(6):1133–46.

9. Khoshkhoo S, Wang Y, Chahine Y, Erson-Omay EZ, Robert SM, Kiziltug E, et al. Contribution of Somatic Ras/Raf/Mitogen-Activated Protein Kinase Variants in the Hippocampus in Drug-Resistant Mesial Temporal Lobe Epilepsy. JAMA Neurol. 2023;80(6):578–87.

10. Mohamed A, Wyllie E, Ruggieri P, Kotagal P, Babb T, Hilbig A, et al. Temporal lobe epilepsy due to hippocampal sclerosis in pediatric candidates for epilepsy surgery. Neurology. 2001;56(12):1643–9.

11. Grunewald R. Childhood seizures and their consequences for the hippocampus. Brain. 2002;125(Pt 9):1935–6.

12. Tartaglia M, Mehler EL, Goldberg R, Zampino G, Brunner HG, Kremer H, et al. Mutations in PTPN11, encoding the protein tyrosine phosphatase SHP-2, cause Noonan syndrome. Nat Genet. 2001;29(4):465–8.

13. Ehrman LA, Nardini D, Ehrman S, Rizvi TA, Gulick J, Krenz M, et al. The protein tyrosine phosphatase Shp2 is required for the generation of oligodendrocyte progenitor cells and myelination in the mouse telencephalon. J Neurosci. 2014;34(10):3767–78.

14. Gauthier AS, Furstoss O, Araki T, Chan R, Neel BG, Kaplan DR, et al. Control of CNS cell-fate decisions by SHP-2 and its dysregulation in Noonan syndrome. Neuron. 2007;54(2):245–62.

15. Glebov-McCloud AGP, Saide WS, Gaine ME, Strack S. Protein Kinase A in neurological disorders. J Neurodev Disord. 2024;16(1):9.

16. Yu ZH, Zhang RY, Walls CD, Chen L, Zhang S, Wu L, et al. Molecular basis of gain-of-function LEOPARD syndrome-associated SHP2 mutations. Biochemistry. 2014;53(25):4136–51.

17. Lauriol J, Kontaridis MI. PTPN11-associated mutations in the heart: has LEOPARD changed Its RASpots? Trends Cardiovasc Med. 2011;21(4):97–104.

18. Tartaglia M, Martinelli S, Stella L, Bocchinfuso G, Flex E, Cordeddu V, et al. Diversity and functional consequences of germline and somatic PTPN11 mutations in human disease. Am J Hum Genet. 2006;78(2):279–90.

19. Johnson H, Ali A, Zhang X, Wang T, Simoulis A, Wingren AG, et al. K-RAS Associated Gene-Mutation-Based Algorithm for Prediction of Treatment Response of Patients with Subtypes of Breast Cancer and Especially Triple-Negative Cancer. Cancers (Basel). 2022;14(21).

20. Prior IA, Lewis PD, Mattos C. A comprehensive survey of Ras mutations in cancer. Cancer Res. 2012;72(10):2457–67.

21. Rabara D, Tran TH, Dharmaiah S, Stephens RM, McCormick F, Simanshu DK, et al. KRAS G13D sensitivity to neurofibromin-mediated GTP hydrolysis. Proc Natl Acad Sci U S A. 2019;116(44):22122–31.

22. Hobbs GA, Der CJ. RAS Mutations Are Not Created Equal. Cancer Discov. 2019;9(6):696–8.

23. Janakiraman M, Vakiani E, Zeng Z, Pratilas CA, Taylor BS, Chitale D, et al. Genomic and biological characterization of exon 4 KRAS mutations in human cancer. Cancer Res. 2010;70(14):5901–11.

24. Schubbert S, Bollag G, Lyubynska N, Nguyen H, Kratz CP, Zenker M, et al. Biochemical and functional characterization of germ line KRAS mutations. Mol Cell Biol. 2007;27(22):7765–70.

25. Bermudez O, Pages G, Gimond C. The dual-specificity MAP kinase phosphatases: critical roles in development and cancer. Am J Physiol Cell Physiol. 2010;299(2):C189–202.

26. Huang M, Wang J, Liu W, Zhou H. Advances in the role of the GADD45 family in neurodevelopmental, neurodegenerative, and neuropsychiatric disorders. Front Neurosci. 2024;18:1349409.

27. Liebermann DA, Hoffman B. Gadd45 in stress signaling. J Mol Signal. 2008;3:15.

28. Liebermann DA, Tront JS, Sha X, Mukherjee K, Mohamed-Hadley A, Hoffman B. Gadd45 stress sensors in malignancy and leukemia. Crit Rev Oncog. 2011;16(1-2):129–40.

29. Low HB, Zhang Y. Regulatory Roles of MAPK Phosphatases in Cancer. Immune Netw. 2016;16(2):85–98.

30. Tubita A, Papini D, Tusa I, Rovida E. Dual-Specificity Protein Phosphatases Targeting Extracellular Signal-Regulated Kinases: Friends or Foes in the Biology of Cancer? Int J Mol Sci. 2025;26(17).

31. Altmuller F, Pothula S, Annamneedi A, Nakhaei-Rad S, Montenegro-Venegas C, Pina-Fernandez E, et al. Aberrant neuronal activity-induced signaling and gene expression in a mouse model of RASopathy. PLoS Genet. 2017;13(3):e1006684.

32. Cantallops I, Routtenberg A. Kainic acid induction of mossy fiber sprouting: dependence on mouse strain. Hippocampus. 2000;10(3):269–73.

33. Faherty CJ, Xanthoudakis S, Smeyne RJ. Caspase-3-dependent neuronal death in the hippocampus following kainic acid treatment. Brain Res Mol Brain Res. 1999;70(1):159–63.

34. Ferraro TN, Golden GT, Smith GG, Schork NJ, St Jean P, Ballas C, et al. Mapping murine loci for seizure response to kainic acid. Mamm Genome. 1997;8(3):200–8.

35. Hu RQ, Koh S, Torgerson T, Cole AJ. Neuronal stress and injury in C57/BL mice after systemic kainic acid administration. Brain Res. 1998;810(1-2):229–40.

36. McKhann GM, 2nd, Wenzel HJ, Robbins CA, Sosunov AA, Schwartzkroin PA. Mouse strain differences in kainic acid sensitivity, seizure behavior, mortality, and hippocampal pathology. Neuroscience. 2003;122(2):551–61.

37. Schauwecker PE, Williams RW, Santos JB. Genetic control of sensitivity to hippocampal cell death induced by kainic acid: a quantitative trait loci analysis. J Comp Neurol. 2004;477(1):96–107.

38. Albala BJ, Moshe SL, Okada R. Kainic-acid-induced seizures: a developmental study. Brain Res. 1984;315(1):139–48.

39. Nickels KC, Wong-Kisiel LC, Moseley BD, Wirrell EC. Temporal lobe epilepsy in children. Epilepsy Res Treat. 2012;2012:849540.

40. Vernet O, Farmer JP, Montes JL, Villemure JG, Meagher-Villemure K. Dysgenetic mesial temporal sclerosis: an unrecognized entity. Childs Nerv Syst. 2000;16(10-11):719–23.

41. Lewis DV, Shinnar S, Hesdorffer DC, Bagiella E, Bello JA, Chan S, et al. Hippocampal sclerosis after febrile status epilepticus: the FEBSTAT study. Ann Neurol. 2014;75(2):178–85.

42. Fabera P, Krijtova H, Tomasek M, Krysl D, Zamecnik J, Mohapl M, et al. Familial temporal lobe epilepsy due to focal cortical dysplasia type IIIa. Seizure. 2015;31:120–3.

43. Blumcke I, Spreafico R. Cause matters: a neuropathological challenge to human epilepsies. Brain Pathol. 2012;22(3):347–9.

44. Barba C, Cossu M, Guerrini R, Di Gennaro G, Villani F, De Palma L, et al. Temporal lobe epilepsy surgery in children and adults: A multicenter study. Epilepsia. 2021;62(1):128–42.

45. Blumcke I, Thom M, Aronica E, Armstrong DD, Bartolomei F, Bernasconi A, et al. International consensus classification of hippocampal sclerosis in temporal lobe epilepsy: a Task Force report from the ILAE Commission on Diagnostic Methods. Epilepsia. 2013;54(7):1315–29.

46. Duchowny M, Levin B, Jayakar P, Resnick T, Alvarez L, Morrison G, et al. Temporal lobectomy in early childhood. Epilepsia. 1992;33(2):298–303.

47. Gunbey C, Soylemezoglu F, Bilginer B, Karli Oguz K, Akalan N, Topcu M, et al. International consensus classification of hippocampal sclerosis and etiologic diversity in children with temporal lobectomy. Epilepsy Behav. 2020;112:107380.

48. McLellan A, Davies S, Heyman I, Harding B, Harkness W, Taylor D, et al. Psychopathology in children with epilepsy before and after temporal lobe resection. Dev Med Child Neurol. 2005;47(10):666–72.

49. Terra-Bustamante VC, Coimbra ER, Rezek KO, Escorsi-Rosset SR, Guarnieri R, Dalmagro CL, et al. Cognitive performance of patients with mesial temporal lobe epilepsy and incidental calcified neurocysticercosis. J Neurol Neurosurg Psychiatry. 2005;76(8):1080–3.

50. Maton B, Jayakar P, Resnick T, Morrison G, Ragheb J, Duchowny M. Surgery for medically intractable temporal lobe epilepsy during early life. Epilepsia. 2008;49(1):80–7.

51. Khoshkhoo S, Bae M, Wang Y, Tillett A, Ramirez RB, Finander B, et al. Activating Ras-MAPK pathway variants drive hippocampal clonal competition in human epilepsy. bioRxiv. 2026.

52. Che Y, Yu YM, Han PL, Lee JK. Delayed induction of p38 MAPKs in reactive astrocytes in the brain of mice after KA-induced seizure. Brain Res Mol Brain Res. 2001;94(1-2):157–65.

53. Hu JH, Malloy C, Hoffman DA. P38 Regulates Kainic Acid-Induced Seizure and Neuronal Firing via Kv4.2 Phosphorylation. Int J Mol Sci. 2020;21(16).

54. Kim SW, Yu YM, Piao CS, Kim JB, Lee JK. Inhibition of delayed induction of p38 mitogen-activated protein kinase attenuates kainic acid-induced neuronal loss in the hippocampus. Brain Res. 2004;1007(1-2):188–91.

55. Namiki K, Nakamura A, Furuya M, Mizuhashi S, Matsuo Y, Tokuhara N, et al. Involvement of p38alpha in kainate-induced seizure and neuronal cell damage. J Recept Signal Transduct Res. 2007;27(2-3):99–111.

56. Poddar R, Paul S. Novel crosstalk between ERK MAPK and p38 MAPK leads to homocysteine-NMDA receptor-mediated neuronal cell death. J Neurochem. 2013;124(4):558–70.

57. Kawataki S, Kubota Y, Katayama K, Imoto S, Takekawa M. GADD45beta-MTK1 signaling axis mediates oncogenic stress-induced activation of the p38 and JNK pathways. Cancer Sci. 2025;116(1):128–42.

58. Kelly BJ, Fitch JR, Hu Y, Corsmeier DJ, Zhong H, Wetzel AN, et al. Churchill: an ultra-fast, deterministic, highly scalable and balanced parallelization strategy for the discovery of human genetic variation in clinical and population-scale genomics. Genome Biol. 2015;16(1):6.

59. Pedersen BS, Quinlan AR. Mosdepth: quick coverage calculation for genomes and exomes. Bioinformatics. 2018;34(5):867–8.

60. Jun G, Flickinger M, Hetrick KN, Romm JM, Doheny KF, Abecasis GR, et al. Detecting and estimating contamination of human DNA samples in sequencing and array-based genotype data. Am J Hum Genet. 2012;91(5):839–48.

61. Cibulskis K, Lawrence MS, Carter SL, Sivachenko A, Jaffe D, Sougnez C, et al. Sensitive detection of somatic point mutations in impure and heterogeneous cancer samples. Nat Biotechnol. 2013;31(3):213–9.

62. Koboldt DC, Zhang Q, Larson DE, Shen D, McLellan MD, Lin L, et al. VarScan 2: somatic mutation and copy number alteration discovery in cancer by exome sequencing. Genome Res. 2012;22(3):568–76.

63. Koboldt DC, Miller KE, Miller AR, Bush JM, McGrath S, Leraas K, et al. PTEN somatic mutations contribute to spectrum of cerebral overgrowth. Brain. 2021;144(10):2971–8.

64. Mirdita M, Schutze K, Moriwaki Y, Heo L, Ovchinnikov S, Steinegger M. ColabFold: making protein folding accessible to all. Nat Methods. 2022;19(6):679–82.

65. Emsley P, Lohkamp B, Scott WG, Cowtan K. Features and development of Coot. Acta Crystallogr D Biol Crystallogr. 2010;66(Pt 4):486–501.

66. Hoffman GE, Roussos P. Dream: powerful differential expression analysis for repeated measures designs. Bioinformatics. 2021;37(2):192–201.

67. Chan G, Kalaitzidis D, Usenko T, Kutok JL, Yang W, Mohi MG, et al. Leukemogenic Ptpn11 causes fatal myeloproliferative disorder via cell-autonomous effects on multiple stages of hematopoiesis. Blood. 2009;113(18):4414–24.

