## Supplemental Table 2 for "Somatic variants activating the RAS-MAPK pathway confer susceptibility to hippocampal sclerosis in drug-resistant epilepsy"

**Supplemental Table 2: Samples and Somatic Findings**

|  | Brain region | Frozen/FFPE | | Neuropathology (detailed) | Somatic Variant | | | VAF % (WES) | | VAF % (targeted) | | Germline  Findings | |
| --- | --- | --- | --- | --- | --- | --- | --- | --- | --- | --- | --- | --- | --- |
|  | 2 - Case - FCD IIId - History of TBI | | | | | | | | | | | | |
|  | Frontal lobe 1 | Frozen | | Neuronal loss and gliosis; white matter gliosis and rarefaction | None | | | - | | - | | - | |
|  | Perisylvian gyrus | Frozen | | Neuronal loss and gliosis; neuronal dyslamination; white matter gliosis and rarefaction | None | | | - | | - | | - | |
|  | Lateral temporal lobe | Frozen | | Neuronal loss and gliosis; white matter gliosis and rarefaction | None | | | - | | - | | - | |
|  | Hippocampus | Frozen | | Hippocampal sclerosis type 1 | None | | | - | | - | | - | |
|  | Prefrontal gyrus | Frozen | | Neuronal loss and gliosis; neuronal dyslamination; white matter gliosis and rarefaction | None | | | - | | - | | - | |
|  | Frontal lobe 2 | Frozen | | Neuronal loss and gliosis; neuronal dyslamination; white matter gliosis and rarefaction | None | | | - | | - | | - | |
|  | Medial frontal lobe | Frozen | | Neuronal loss and gliosis; neuronal dyslamination; white matter gliosis and rarefaction | None | | | - | | - | | - | |
|  | Temporal occipital lobe | Frozen | | Neuronal loss and gliosis; neuronal dyslamination; white matter gliosis and rarefaction | None | | | - | | - | | - | |
|  | Occipital lobe | Frozen | | Neuronal loss and gliosis; neuronal dyslamination; white matter gliosis and rarefaction | None | | | - | | - | | - | |
|  | Parietal lobe | Frozen | | Neuronal loss and gliosis; neuronal dyslamination; white matter gliosis and rarefaction | None | | | - | | - | | - | |
|  | 8 - Case - FCD IIId - History of perinatal stroke | | | | | | | | | | | | |
|  | Frontal lobe | Frozen | | Encephaloclastic changes; possible dyslamination | None | | | - | | - | | - | |
|  | Medial frontal lobe | Frozen | | Encephaloclastic changes | None | | | - | | - | | - | |
|  | Posterior frontal lobe | Frozen | | Encephaloclastic changes; possible dyslamination | None | | | - | | - | | - | |
|  | Posterior lateral frontal lobe | Frozen | | Encephaloclastic changes; possible dyslamination | None | | | - | | - | | - | |
|  | Parietal lobe | Frozen | | Encephaloclastic changes; possible dyslamination | None | | | - | | - | | - | |
|  | Occipital lobe | Frozen | | Encephaloclastic changes; possible dyslamination | None | | | - | | - | | - | |
|  | Lateral temporal lobe | Frozen | | Encephaloclastic changes; possible dyslamination | None | | | - | | - | | - | |
|  | Hippocampus | Frozen | | Hippocampal sclerosis type 1 | None | | | - | | - | | - | |
|  | Cingulate gyrus | Frozen | | Encephaloclastic changes | None | | | - | | - | | - | |
|  | 9 - Case - FCD IIIa - History of febrile seizure | | | | | | | | | | | | |
|  | Anterior temporal lobe; lateral neocortex | Frozen | | Cortical dyslamination; heterotopic gray matter in white matter | None | | | - | | - | | NM_001447.2(FAT2):c.10758G>C p.(K3586N) | |
|  | Residual hippocampus | FFPE | | Gliosis | None | | | - | | - | |  |  |
|  | Hippocampal tail | FFPE | | Reactive changes | None | | | - | | - | |  |  |
|  | 13 - Case - FCD IIa with HS Type I - History of febrile status epilepticus | | | | | | | | | | | | |
|  | Frontal operculum | Frozen | | Neuronal dyslamination with dysmorphic neurons | None | | | - | | - | | - | |
|  | Anterior temporal lobe; lateral neocortex | Frozen | | Neuronal dyslamination with dysmorphic neurons | None | | | - | | - | | - | |
|  | Amygdala | FFPE | | Gliosis | None | | | - | | - | | - | |
|  | Hippocampus | FFPE | | Hippocampal sclerosis type 1 | None | | | - | | - | | - | |
|  | Parietal operculum | FFPE | | Neuronal dyslamination with dysmorphic neurons | None | | | - | | - | | - | |
|  | Supramarginal gyrus | FFPE | | Neuronal dyslamination with dysmorphic neurons | None | | | - | | - | | - | |
|  | 36 - Case - FCD IIIa / Diffuse FCD | | | | | | | | | | | | |
|  | Superior temporal gyrus | FFPE | | Probably neuronal dyslamination | NM_002834.5(PTPN11):c.1502G>A p.(R501K) | | | NS | | 0 | | - | |
|  | Amygdala | FFPE | | Gliosis |  |  |  | NS | | 0 | | - | |
|  | Hippocampus | FFPE | | Hippocampal sclerosis |  |  |  | NS | | 0.22 | | - | |
|  | Inferior parietal | Frozen | | Cortical dyslamination |  |  |  | 0 | | 0 | | - | |
|  | Posterior inferior frontal | Frozen | | Cortical dyslamination |  |  |  | 2.96 | | 3.32 | | - | |
|  | 38 - Case - FCD Ic with HS Type I | | | | | | | | | | | | |
|  | Inferior temporal lobe | Frozen | | Cortical dyslamination | None | | | - | | - | | NM_001100913.3(PACS2):c.2147del p.(G716fs) | |
|  | Posterior temporal lobe | Frozen | | Cortical dyslamination | None | | | - | | - | |  |  |
|  | Amygdala | Frozen | | Gliosis | None | | | - | | - | |  |  |
|  | Inferior parietal cortex | Frozen | | Cortical dyslamination | None | | | - | | - | |  |  |
|  | Posterior temporal lobe | Frozen | | Cortical dyslamination | None | | | - | | - | |  |  |
|  | Hippocampus | FFPE | | Hippocampal sclerosis with prominent neuronal loss in CA1 / CA4 | None | | | - | | - | |  |  |
|  | 58 - Case - FCD IIa with HS - History of febrile seizure | | | | | | | | | | | | |
|  | Lateral temporal lobe | Frozen | | Cortical dyslamination with reactive changes | None | | | - | | - | | - | |
|  | Posterior temporal lobe | Frozen | | Cortical dyslamination with reactive changes | None | | | - | | - | | - | |
|  | Frontal operculum | Frozen | | Cortical dyslamination with reactive changes | None | | | - | | - | | - | |
|  | Parietal operculum | Frozen | | Cortical dyslamination with reactive changes | None | | | - | | - | | - | |
|  | Amygdala | Frozen | | Gliosis and possible heterotopia/hamartoma | None | | | - | | - | | - | |
|  | Hippocampus | Frozen | | Hippocampal sclerosis with prominent neuronal loss in CA1 / CA4 | None | | | - | | - | | - | |
|  | 81 - Case - FCD IIb with HS | | | | | | | | | | | | |
|  | Temporal lobe lesion | Frozen | | Focus of dysmorphic neurons and microcalcifications | None | | | - | | - | | - | |
|  | Anterior lateral temporal lobe | Frozen | | Cortical dyslamination, focal nodular gray matter heterotopia | None | | | - | | - | | - | |
|  | Posterior temporal lobe | Frozen | | Cortical dyslamination with dysmorphic neurons and balloon cells | None | | | - | | - | | - | |
|  | Hippocampus | Frozen | | Neuronal loss and gliosis | None | | | - | | - | | - | |
|  | Amygdala | Frozen | | Cortical dyslamination | None | | | - | | - | | - | |
|  | 85 - Case - FCD IIIa | | | | | | | | | | | | |
|  | Anterior temporal lobe; lateral neocortex | Frozen | | Cortical dyslamination | None | | | - | | - | | - | |
|  | Superior temporal gyrus | NS | | Gliosis | None | | | - | | - | | - | |
|  | Amygdala | NS | | Gliosis | None | | | - | | - | | - | |
|  | Hippocampal head | Frozen | | Gliosis and neuronal loss | None | | | - | | - | | - | |
|  | 86 - Case - FCD IIIa - History of status epilepticus | | | | | | | | | | | | |
|  | Lateral temporal lobe | Frozen | | Patchy cortical and diffuse white matter gliosis | *PRKACB*  12 kb deletion; exons 2 - 5 | | | NA | | - | | - | |
|  | Hippocampal head | Frozen | | Gliosis and neuronal dropout |  |  |  | NA | | - | | - | |
|  | 107 - Case - FCD IIIa - History of febrile seizure | | | | | | | | | | | | |
|  | Lateral temporal cortex | Frozen | | Cortical dyslamination | NM_002834.5(PTPN11):c.1505C>T p.(S502L) | | | NS | | 0 | | - | |
|  | Hippocampus | Frozen | | Hippocampal sclerosis with severe neuronal loss in CA1 and dentate duplication |  |  |  | 4.10 | | 3.80 | | - | |
|  | 109 - Case - FCD IIIb / IIIa Dual Pathology | | | | | | | | | | | | |
|  | Posterior temporal lobe | Frozen | | Mild cortical dyslamination and gliosis; hypercellular white matter and gliosis | NM_004333.6 (BRAF): c.1799T>A p.(V600E) | | | 0.56 | | 0 | | - | |
|  | Anterior temporal tip | FFPE | | Focal cortical dysplasia; gliosis; microcalcification |  |  |  | 4.48 | | 0 | | - | |
|  | Hippocampus | FFPE | | Mesial temporal / hippocampal sclerosis type 1b; low-grade glioneuronal lesion |  |  |  | 23.84 | | 19.41 | | - | |
|  | Superior temporal gyrus | FFPE | | Low-grade glioneuronal lesion |  |  |  | NS | | - | | - | |
|  | 121 - Case - FCD 1c and MTS - History of status epilepticus | | | | | | | | | | | | |
|  | Lateral neocortex | Frozen | | Cortical dyslamination with diffuse gliosis | NM_004985.5(KRAS):c.350A>G p.(K117R) | | | 7.99 | | 12.47 | | - | |
|  | Superior temporal gyrus | Frozen | | Fragment of cortex and underlying white matter with gliosis |  |  |  | 2.61 | | 3.95 | | - | |
|  | 122 - Case - FCD IIIa | | | | | | | | | | | | |
|  | Anterior temporal lobe; lateral neocortex | Frozen | | Cortical dyslamination; diffuse cortical and white matter gliosis | NM_002834.5(PTPN11):  c.1381G>A  p.(A461T) | | | 0 | | 0 | | - | |
|  | Amygdala | Frozen | | Neuronal dropout and diffuse gliosis |  |  |  | 0 | | 0 | | - | |
|  | Hippocampus head and body | Frozen | | Neuronal dropout (CA1 predominant) and diffuse gliosis |  |  |  | 5.97 | | 12.78 | | - | |
|  | Parahippocampal gyrus | Frozen | | Cortical dyslamination and diffuse gliosis |  |  |  | 1.8 | | 2.07 | | - | |
|  | Hippocampal tail | Frozen | | Diffuse gliosis |  |  |  | NS | | NS | | - | |
|  | 129 - Case - FCD IIIa - History of status epilepticus | | | | | | | | | | | | |
|  | Lateral neocortex | Frozen | | Mild cortical dyslamination | None | | | - | | - | | - | |
|  | Hippocampus | Frozen | | Focal necrosis; diffuse gliosis and reactive changes | None | | | - | | - | | - | |
|  | 1 - Control - FCD Ic - History of Cryptogenic West Syndrome | | | | | | | | | | | | |
|  | Left temporal lobe; lateral neocortex | Frozen | | Cortical dyslamination | NM_005660.2(SLC35A2):c.634_635delTC p.(S212Lfs) | | 2.96 | | | 2.67 | | - | |
|  | Left Amygdala | Frozen | | Fragment of gray and white matter with gliosis |  |  | 12.5 | | | 5.2 | | - | |
|  | Left Hippocampus; head and body | Frozen | | Fragment of gray and white matter with gliosis |  |  | 27.67 | | | 13.51 | | - | |
|  | Left occipital lobe and pole | Frozen | | Cortical dyslamination |  |  | 9.8 | | | 2.84 | | - | |
|  | Portion of left superior temporal gyrus | FFPE | | Cortical dyslamination |  |  | - | | | 5.25 | | - | |
|  | Left inferior occipital lobe | FFPE | | Cortical dyslamination |  |  | - | | | 1.66 | | - | |
|  | Left medial occipital lobe | FFPE | | Cortical dyslamination |  |  | - | | | 11.81 | | - | |
|  | Left inferior parietal cortex | FFPE | | Cortical dyslamination |  |  | - | | | 8.67 | | - | |
|  | 16 - Control - FCD Ic - History of focal seizures and infantile spasms | | | | | | | | | | | | |
|  | Right frontal cortex | FFPE | | NA - 2013 surgery | chr 1q gain | | | 10.39 | | - | | - | |
|  | Right frontal operculum | Frozen | | Cortical dyslamination with reactive changes |  |  |  | 5.31 | | - | | - | |
|  | Right inferior posterior frontal gyrus | Frozen | | Cortex and white matter with reactive changes |  |  |  | 13.73 | | - | | - | |
|  | Right temporal lobe, lateral cortex | Frozen | | Cortical dyslamination |  |  |  | 0.50 | | - | | - | |
|  | Right amygdala | Frozen | | Fragment of gray and white matter with gliosis |  |  |  | 1.25 | | - | | - | |
|  | Right hippocampus | Frozen | | hippocampus with gliosis |  |  |  | 0.14 | | - | | - | |
|  | 27 - Control - FCD IIId - History of Perinatal right middle cerebral artery infarct | | | | | | | | | | | | |
|  | Right temporal lobe, lateral neocortex | | Frozen | Focal cortical dyslamination and white matter gliosis | | None | | | - | | - | | **-** |
|  | Right hippocampus | | Frozen | hippocampal and white matter gliosis | | None | | | - | | - | | **-** |
|  | 28 - Control - FCD IIIa - Possible history of encephalitis / meningitis and febrile seizures | | | | | | | | | | | | |
|  | Inferior lateral temporal lobe | Frozen | | Cortical dyslamination with white matter gliosis | None | | | - | | - | | - | |
|  | Left frontal operculum | Frozen | | Cortical dyslamination; focal superficial glial scar and white matter gliosis | None | | | - | | - | | - | |
|  | Left posterior temporal lobe | Frozen | | Cortical dyslamination with white matter gliosis | None | | | - | | - | | - | |
|  | Left parietal operculum | Frozen | | Cortical dyslamination with white matter gliosis | None | | | - | | - | | - | |
|  | Amygdala | Frozen | | Gray and white matter with gliosis | None | | | - | | - | | - | |
|  | Hippocampus | Frozen | | hippocampal and white matter gliosis | None | | | - | | - | | - | |
|  | 43 - Control - FCD Ic - History of West syndrome and infantile spasms | | | | | | | | | | | | |
|  | Lateral temporal lobe | Frozen | | Malformation of cortical development, gliosis, reactive astrocytosis | chr 1q gain | | | 3.86 | | - | | - | |
|  | Frontoparietal operculum | Frozen | | Malformation of cortical development, gliosis, reactive astrocytosis |  |  |  | 20.18 | | - | | - | |
|  | Amygdala | Frozen | | gliosis, reactive astrocytosis |  |  |  | 1.02 | | - | | - | |
|  | Parahippocampus | Frozen | | Malformation of cortical development, gliosis, reactive astrocytosis |  |  |  | 0.44 | | - | | - | |
|  | Hippocampus head and body | Frozen | | Malformation of cortical development (in adjacent cortex), gliosis / reactive astrocytosis in adjacent white matter and subpial zone |  |  |  | 0.37 | | - | | - | |
|  | 51 - Control - FCD Ia - History of right temporal focus nonmotor seizures | | | | | | | | | | | | |
|  | Right anterior temporal lobe | Frozen | | focal cortical dysplasia, gliosis | None | | | - | | - | | - | |
|  | Right amygdala | Frozen | | Increased white matter neurons, gliosis | None | | | - | | - | | - | |
|  | Right hippocampal head | Frozen | | Increased white matter neurons, reactive astrocytosis in adjacent white matter | None | | | - | | - | | - | |
|  | 60 - Control - FCD IIId - History of left ventricular thrombus and right middle cerebral artery stroke | | | | | | | | | | | | |
|  | Right inferior frontal gyrus | Frozen | | Cortical dyslamination | None | | | - | | - | | - | |
|  | Right parietal operculum supramarginal gyrus | Frozen | | Cortical dyslamination | None | | | - | | - | | - | |
|  | Right anterior temporal lobe | Frozen | | Cortical dyslamination | None | | | - | | - | | - | |
|  | Right amygdala | Frozen | | Gray and white matter with gliosis | None | | | - | | - | | - | |
|  | Right hippocampus and Parahippocampus lesion | Frozen | | hippocampus with gliosis | None | | | - | | - | | - | |
| 94 - Control - FCD IIId - History of Infantile spasms, encephalomalacia, metopic craniosynostosis, microcephaly | | | | | | | | | | | | | |
|  | Right fronto-parietal operculum | Frozen | | Cortical dyslamination associated with gliomesodermal scar, white matter gliosis | None | | | - | | - | | - | |
|  | Right temporal lobe, lateral neocortex | Frozen | | Cortical dyslamination associated with gliomesodermal scar, white matter gliosis | None | | | - | | - | | - | |
|  | Right amygdala | Frozen | | Gray matter with gliosis and mild neuronal dropout | None | | | - | | - | | - | |
|  | Right hippocampus | Frozen | | hippocampal and white matter gliosis | None | | | - | | - | | - | |
|  | 111 - Control - FCD Ia - History of status epilepticus | | | | | | | | | | | | |
|  | Lateral neocortex | Frozen | | Persistent vertical lamination, gliosis | None | | | - | | - | | - | |
|  | Amygdala | Frozen | | gliosis | None | | | - | | - | | - | |
|  | Hippocampus | Frozen | | vertical lamination, gliosis | None | | | - | | - | | - | |
|  | 117 - Control - Diffuse MCD / heterotopias - History of epileptic encephalopathy | | | | | | | | | | | | |
|  | Left frontal operculum | Frozen | | Cortical dyslamination | None | | | - | | - | | - | |
|  | Left parietal operculum | Frozen | | Cortical dyslamination | None | | | - | | - | | - | |
|  | Left temporal lobe 1 | Frozen | | Cortical dyslamination with gray matter heterotopias in white matter | None | | | - | | - | | - | |
|  | Left temporal lobe 2 | Frozen | | Cortical dyslamination with gray matter heterotopias in white matter | None | | | - | | - | | - | |
|  | Left amygdala | Frozen | | Gray and white matter with gliosis | None | | | - | | - | | - | |
|  | Left hippocampus | Frozen | | Hippocampal tissue with gliosis | None | | | - | | - | | - | |

NS: Not sequenced; NA: not applicable
