## Supplemental Table 5 for "Somatic variants activating the RAS-MAPK pathway confer susceptibility to hippocampal sclerosis in drug-resistant epilepsy"

**Supplemental Table 5: Primers for Targeted Amplicon Sequencing**

| Patient ID | Gene | Forward Primer (5'-->3') | Reverse Primer (5'-->3') |
| --- | --- | --- | --- |
| 36 | PTPN11 | CTTCGTAGGTGTTGACTGCGA | GAGCCTGTCCTCCTGCTCAA |
| 107 | PTPN11 | TCGTAGGTGTTGACTGCGAT | GAATGAGAATCCGCATGCCAG |
| 109 | BRAF | GCAGCATCTCAGGGCCAAAA | GCTTGCTCTGATAGGAAAATGAGA |
| 121 | KRAS | AAGTCCTGAGCCTGTTTTGTGT | TTTATGACAAAAGTTGTGGACAGG |
| 122 | PTPN11 | GCTTTTTGTCCTTCTGCCCG | GCTTCTTGCCCACCAGATGA |
| 1 | SLC35A2 | AGCCTGAGCTGCCTTTG | CGGCCACTGGATCAGAAC |
