## Supplemental Table 1 for "Somatic variants activating the RAS-MAPK pathway confer susceptibility to hippocampal sclerosis in drug-resistant epilepsy"

**Supplemental Table 1: Patients (Cases/Controls)**

| Patient ID | Case/  Control | Sex | Onset Age  (years) | Surgery Age  (years) | Relevant  history | MRI Findings | Surgery | Neuro-pathology (ILAE classification) |
| --- | --- | --- | --- | --- | --- | --- | --- | --- |
| 2 | Case | M | 3 | 11 | Traumatic brain injury | Diffuse injury to left hemisphere with encephalocele | Left anatomical hemispherectomy with pseudo meningeal cyst repair | FCD IIId and HS Type I |
| 8 | Case | F | 4.5 | 12 | Premature birth with hydrocephalus, perinatal stroke | Enlarged ventricles, encephalomalacia, reduced white matter volume, bilateral moderate hippocampal atrophy without increased FLAIR signal | Right anatomical hemispherectomy | FCD IIId and HS Type I |
| 9 | Case | F | 8 | 11 (1^st^ surgery), 12 (2^nd^ surgery) | 30-min febrile seizure at 11 months of age | Findings suggestive of left hippocampal sclerosis | Left amygdalo-hippocampotomy and left temporal lobectomy with amygdalo-hippocampectomy | FCD IIIa |
| 13 | Case | F | 0 | 11 | Infantile spasms; febrile status epilepticus at 3 years of age; Lennox Gastaut secondary to hemiconvulsion-hemiplegia-epilepsy syndrome | Right hemispheric atrophy, periventricular gliosis, basal ganglia atrophy, and right mesial temporal sclerosis (MTS) | Right peri-insular hemispherectomy | FCD IIa and HS Type 1a |
| 36 | Case | F | 0.5 | 1 (1st surgery), 3 (2^nd^ surgery) | None | Signs of volume loss in left hippocampal head suggestive of developing mesial temporal sclerosis. Optic pathway glioma, left cerebral hypoplasia/dysplasia | Left temporal lobectomy and left peri-insular functional hemispherotomy | FCD IIIa / Diffuse FCD |
| 38 | Case | M | 11 | 14 | None | Mild to moderate atrophy of right frontal lobe; moderate atrophy and increased FLAIR signal involving right hippocampal body and head, consistent with mesial temporal sclerosis | Right temporal lobectomy and inferior parietal cortical resection | FCD Ic and HS Type Ia |
| 58 | Case | M | 4 | 13 (1^st^ surgery), 17 (2^nd^ surgery) | Febrile seizure and encephalitis at 4 months of age | Marked volume loss of the right cerebral hemisphere and the left cerebellar hemisphere, atrophic right hippocampus | Corpus callosotomy (1^st^ surgery), right hemisphere excision of lateral temporal lobe, posterior temporal lobe, frontal operculum, parietal operculum, amygdala, and hippocampus (2^nd^ surgery) | FCD IIa and HS |
| 81 | Case | F | 2 | 4 | None | Amorphous calcification at the depth of the posterior aspect of left inferior temporal sulcus suggestive of calcified dysplasia or glioneuronal tumor | Resection of left temporal lesion, angular gyrus, mesial temporal lobe, middle and inferior temporal gyrus | FCD IIb with HS |
| 85 | Case | M | 4 | 6 | None | Subtle blurring of cortical-subcortical margins within left inferior temporal and/or left occipito-temporal (fusiform) regions | Left anterior temporal lobectomy with amygdalo-hippocampectomy | FCD IIIa |
| 86 | Case | F | 8 | 10 | Status epilepticus | Left temporal encephalocele, developing left mesial temporal sclerosis with left hippocampal head volume loss and increased T2-weighted signal | Left temporal lobectomy and repair of the skull left temporal encephalocele | FCD IIIa |
| 107 | Case | M | 3 | 9 | Febrile seizure at 15 months of age | Decreased volume and diffuse FLAIR hyperintensity of the left temporal lobe and hippocampus with decreased gray-white differentiation concerning for left temporal lobe dysplasia, left mesial temporal sclerosis | Left temporal lobectomy | FCD IIIa |
| 109 | Case | M | 2 | 3 (1^st^ surgery), 13 (2^nd^ surgery) | Left temporal ganglioglioma | Areas of abnormality in the left anterior and mesial temporal/hippocampal | Left temporal ganglioglioma resection and left anterior temporal lobectomy with amygdalohippocampectomy (1^st^ surgery), revision of temporal lobectomy (2^nd^ surgery) | FCD IIIb / IIIa Dual Pathology |
| 121 | Case | F | 7 | 11 | Status epilepticus | Moderate atrophy and increased FLAIR signal within left hippocampal formation, consistent with left mesial temporal sclerosis. | Left stereotactic laser amygdalohippocampectomy (1^st^ surgery), Left temporal lobectomy (2^nd^ surgery) | FCD Ic and MTS |
| 122 | Case | M | 4 | 10 | Hemizygous for X-linked OPHN1 germline variant, mild inferior vermian hypoplasia consistent with mild Dandy-Walker sequence abnormalities | Left-sided mesial temporal sclerosis | Left anterior temporal lobectomy with amygdalohippocampectomy | FCD IIIa |
| 129 | Case | M | 16 | 20 | Status epilepticus | Left mesial temporal sclerosis | Ablation of left hippocampus and amygdala (1^st^ surgery), Left temporal lobectomy (2^nd^ surgery) | FCD IIIa |
| 1 | Control | M | 0.75 | 3 | West syndrome | Abnormal increased signal of the left temporal white matter with blurring of the gray-white matter interface | Left Temporo-Parieto-Occipital craniotomy | FCD Ic |
| 16 | Control | F | 0.33 | 3 (1^st^ surgery), 8 (2^nd^ surgery) | Infantile spasms | Stable encephalomalacia involving right superior and inferior parietal lobules and right precentral gyrus | Resection of right frontal FCD (1^st^ surgery), right peri-insular functional hemispherotomy (2^nd^ surgery) | FCD Ic |
| 27 | Control | F | 3 | 5 | Perinatal right middle cerebral artery infarct | Stable chronic encephalomalacia in the distribution of the right middle cerebral artery consistent with old infarct; associated gliosis along the margins of the lesion | Right temporal lobe lateral neocortex resection; right hippocampus resection | FCD IIId |
| 28 | Control | F | unknown | 10 | Febrile seizures and encephalitis / meningitis | Diffuse atrophy of the left hemisphere, T2H in left posterior quadrant | Left hemispherotomy | FCD IIId |
| 43 | Control | M | 0.16 | 1 | West syndrome, infantile spasms | Left frontal polymicrogyria | Left hemispherotomy | FCD Ic |
| 51 | Control | F | 7 | 9 | None | Diminished cerebral flood flow suggested in the anterior right temporal lobe, potentially reflecting an occult right-sided epileptogenic focus | Craniotomy with right anterior temporal lobectomy and amygdala-hippocampectomy | FCD Ia |
| 60 | Control | F | 2 | 4 | Left ventricular thrombus and right middle cerebral artery stroke | Encephalomalacia; remote infarct | Excision of right inferior frontal gyrus, parietal operculum supramarginal gyrus, right anterior temporal lobe lateral neocortex, right amygdala, right hippocampus. | FCD IIId |
| 94 | Control | M | 0 | 2 | Infantile spasms; encephalomalacia, metopic craniosynostosis, microcephaly | Severe encephalomalacia, ex vacuo dilation of ventricles, thinning corpus callosum | Right hemispherectomy | FCD IIId |
| 111 | Control | M | 4 | 14 | Status epilepticus | Enlargement of right amygdala and mild T2 hyperintense signal, small left hippocampal head without signal abnormality. | Right craniotomy with right temporal lobectomy | FCD Ia |
| 117 | Control | M | 0.75 | 2 | Epileptic encephalopathy | Left-sided frontal-temporal-parietal open lip schizencephaly with a wide cleft; small left cerebral hemisphere with thin corpus callosum, hypoplastic left basal ganglia and thalamus; multiple foci of heterotopic gray matter along the left lateral ventricle | Left peri-insular functional hemispherotomy | Diffuse FCD |
